# A new molecular seed assay to predict *Ustilago nuda* field infection levels

**DOI:** 10.1101/2025.06.25.661478

**Authors:** Cecilia Panzetti, Eveline Jenny, Irene Bänziger, Andreas Kägi, Susanne Vogelgsang, Thomas Hebeisen, Peter Büttner, Daniel Croll, Franco Widmer, Karen E. Sullam

## Abstract

Seed health tests are performed to prevent sowing untreated seeds with problematic pathogen levels. The detection of internal seedborne pathogens like *Ustilago nuda*, causing loose smut in barley (*Hordeum vulgare*), is challenging because symptoms only appear when teliospores replace barley inflorescences and smutted ears develop. Currently, *U. nuda* seed infection levels are determined from the visual inspections of extracted embryos or fields of plants used for seed production, both of which are laborious and can be unreliable. To improve *U. nuda* detection, we developed a multiplex qPCR method targeting *U. nuda* and *H. vulgare* DNA. Naturally infected seed lots were tested using this qPCR method and the visual analysis of embryos. We grew the same seed lots in the field over two seasons and used the observed smutted ears as the reference infection level. The qPCR results, *U. nuda* DNA normalized to *H. vulgare* DNA, better correlated with observed field infections than the number of infected embryos. Our qPCR method more accurately distinguished seed lots with infections above and below the field tolerance threshold. It offers a reliable alternative to the visual analysis of extracted embryos. The integration of our qPCR method with field observations can enhance *U. nuda* management and reduce unnecessary prophylactic seed treatments, thereby bolstering integrated pest management strategies.

## Introduction

Seed health tests detect seedborne pathogens and can serve a foundational role in integrative pest management (IPM) strategies. Plant pathogen detection on untreated seed and the subsequent avoidance of infected material can prevent the spread of seedborne diseases. Indeed, a core IPM principle is the targeted use of synthetic or organic plant protection products (PPPs) only when a harmful organism’s level exceeds a tolerance threshold^1^. Seed health tests detect pathogens, carried on or within seeds^2^, and help determine when these tolerance thresholds are reached and whether seed treatments are necessary^3^. However, often seed suppliers routinely treat seeds with synthetic PPPs by default^3,4^, regardless of the pathogen’s level. Seed treatments are seen as an inexpensive, low effort method to protect crops^5^. As more policies aim to reduce synthetic PPP applications^1^, an increased reliance on seed health tests can reduce unnecessary seed treatments. This targeted approach would ensure that PPPs are applied only when needed based on a predefined tolerance threshold to prevent the resurgence of seedborne diseases^6–9^.

Seedborne pathogens, unlike infections in other plant tissues, can be asymptomatic and difficult to detect in seeds. Undetected pathogens can develop within healthy-appearing plants and then manifest disease symptoms at different plant growth stages^10,11^. Internally located fungal seed pathogens, which can be asymptomatic, are a particular threat to agricultural production because they are difficult to detect and can spread unnoticed. For example, *Ustilago nuda* and *Ustilago tritici*, which cause loose smut in barley (*Hordeum vulgare*) and wheat (*Triticum aestivum*) respectively, can cause significant yield losses^12^. In these diseases, the symptoms become visible when the fungus replaces the plant inflorescence with its teliospores, causing smutted ears, and spreads to nearby healthy plants^10,13,14^. Because these pathogens are asymptomatic through plant growth until ear development, early detection and control remain challenging^15^. Due to their difficulty in early detection, these seedborne pathogens confer a high risk of disease transmission in seed production^16,17^. Since crop losses from *U. nuda* can reach up to 25% in barley and only up to 5% in wheat due to *U. tritici*^18^, loose smut of barley is typically more problematic in cereal seed production.

In barley seed production, as part of the seed certification process, field inspections are conducted to identify diseases. Loose smut infections are assessed during the flowering stage to minimize its propagation to the next generation of barley seed^19^. In Switzerland, for example, no more than 2 infected ears per 100 *m*^2^ are allowed in plants that produce first-generation certified seed and up to 5 infected ears per 100 *m*^2^ in plants that produce second-generation certified seeds that are commercially available^20–22^, but the tolerance thresholds are country-dependent. However, infection detected during the field inspection may not accurately correlate with infection levels in the subsequently harvested seed; environmental factors, such as wind and rain, can hinder the visual detectability of infected ears and increase teliospores dispersion^23^. Although an inspected field may meet the seed certification criterion regarding loose smut, its barley inflorescences may hide *U. nuda* latent infections due to teliospore dispersal from a neighboring field. An unexpectedly high infection rate in the harvested seed may occur despite the appearance of healthy-looking plants^16,17,19,23^. Therefore, weather conditions and teliospore dispersion may obscure the actual number of diseased ears and may lead to an underestimation of the harvested seed’s true infection rate^23^.

A direct assessment of the harvested seeds can avert a potential mismatch between the harvested seed’s *U. nuda* infection rate and the visible field infections during seed production. Although not required for seed certification in many countries^20^, the *U. nuda* infection rate can be determined directly from the inspections of harvested seed rather than an inference based on observed smutted ears in the field during seed production. A validated seed testing protocol for *U. nuda* detection in embryos of barley seed is used in some countries either for seed health certification or as a complement to it^7,20,22,24^. The percentage of infected embryos tolerated for seed health certification depends on the country and ranges from 0.1% to 1.0%^20,22^. Nevertheless, the detection of *U. nuda* mycelia within embryos remains challenging due to limited mycelial visibility, constraints on sample size, and the labor-intensive nature of the process^24–26^.

Molecular methods can improve the detection of seedborne pathogens because such methods increase scalability, sensitivity, and specificity compared to visual inspections^27,28^. Detection protocols for other seedborne fungal pathogens have employed molecular methods successfully^29–32^ and have shown promising results for detecting *U. nuda* in plant material^10,33–35^. Previous research on *U. nuda* detection employed enzyme-linked immunosorbent assay (ELISA) and polymerase chain reaction (PCR) methods^10,33–35^. Among the published techniques that focused on *U. nuda* detection in seed^33,34^, seeds were individually tested, which is neither economical nor feasible for a large-scale application. To facilitate an increased throughput, the bulk analysis of milled seeds can simplify seed sample preparation for PCR-based assays^34,35^. However, initial attempts at using PCR with bulk milled seed samples encountered challenges; in particular, nonspecific fluorescence in SYBR Green-based quantitative PCR (qPCR) assays impaired at the detectability of low *U. nuda* infection levels^35^. This nonspecific fluorescence was observed with a previously published *U. nuda* primer set used on seedling^10^. Within a later publication, this primer set was modified with the addition of three base pairs, and it was applied to seed using a SYBR Green-based qPCR assays^36^. A drawback to this published protocol is that SYBR Green dye binds to all double stranded products and can result in decreased specificity. Therefore, a fluorogenic probe can enhance *U. nuda* assay specificity by providing an additional specific-binding region of the amplified product required for a florescent signal’s production^35^.

No established molecular method can reliably detect *U. nuda* in bulk milled seed samples with the specificity and sensitivity required for seed health testing. In this study, we developed a multiplex qPCR method using newly designed primers and fluorogenic probes that quantify *U. nuda* and *H. vulgare* DNA in bulk milled seed. The host DNA acts as an internal control for the DNA extractions and qPCR reactions. It also normalizes the pathogen DNA in each sample. Furthermore, no prior study has directly connected *U. nuda* seed infection rates detected with molecular methods to infection rates observed in the field. This link is needed to translate the qPCR results directly to field infection levels and, thus, potential yield losses. We used field observations as a reference to evaluate our qPCR detection method’s performance. Additionally, we evaluated the performance of the embryo test —the recognized seed health test for *U. nuda*^24^ —on the same seed lots. Compared to the embryo test, the qPCR method more accurately distinguished seed lots with infection above or below its tolerance threshold, reflecting the classification of the observed field infection levels with respect to the field tolerance threshold. Overall, our newly developed qPCR method showed an improved ability to detect *U. nuda* and predict field infections compared to the embryo test. Due to its increased scalability compared to the embryo test, our qPCR method is also highly applicable for large-scale seed health assessments. It provides a quantitative evaluation of *U. nuda* infections, which enables informed seed management decisions based on a tolerance threshold. Our qPCR method’s incorporation into the barley seed certification process would strengthen IPM strategies to limit loose smut’s spread and reduce the reliance on PPP seed treatments.

## Results

### Sensitive and specific primers and probes

We developed a new multiplex TaqMan qPCR protocol that targets COX3 in the pathogen, *Ustilago nuda*, and GADPH in the host, *Hordeum vulgare*, enabling the normalization of pathogen to host DNA. Our protocol’s limit of detection (LOD) for the *U. nuda* COX3 gene is 80 pg (4 copies per reaction) based on a dilution series of its gBlock gene fragment tested eight times. We also compared our *U. nuda* primers in a multiplex TaqMan reaction with the most recently published ITS primers in a SYBR Green singleplex reaction^36^. Both protocols successfully amplified *U. nuda* teliospores in reactions with template DNA concentrations of 0.8, 0.08, and 0.008 ng/µL (Table 1). From our tests to determine the primer specificity, we observed that our COX3 primers also amplified *Ustilago hordei* mycelium, which was expected from the sequence similarities of *U. nuda* and *U. hordei* at the primer binding sites (Supplementary Information, Figure 1). We tested DNA from mycelial samples of other basidiomycetes that had been isolated from single teliospores to determine the COX3 and ITS primers’ specificity. The published ITS protocol consistently amplified all *Ustilago* species tested with the DNA template concentration of 0.8 ng/µL per reaction (Table 1). In contrast, our newly developed protocol amplified *Ustilago maydis* in one of three replicates with a quantification cycle (Cq) of 36.42, and it did not amplify another non-target *Ustilago* spp., showing increased specificity. Additionally, the ITS protocol amplified an unidentified *Pseudozyma* species (Cq = 37.51 *±*0.36), while our protocol showed no amplification. The other six non-target fungal taxa exhibited either unstable amplification (i.e. amplification in only some replicates) or no amplification in both protocols (Table 1). We found that the coefficient of variation (CV) was 89% lower with our protocol than with the ITS protocol when applied to *U. nuda* infected seed and seedling samples (Supplementary Information, Table 1), showing in more stable measurements.

**Table 1.**
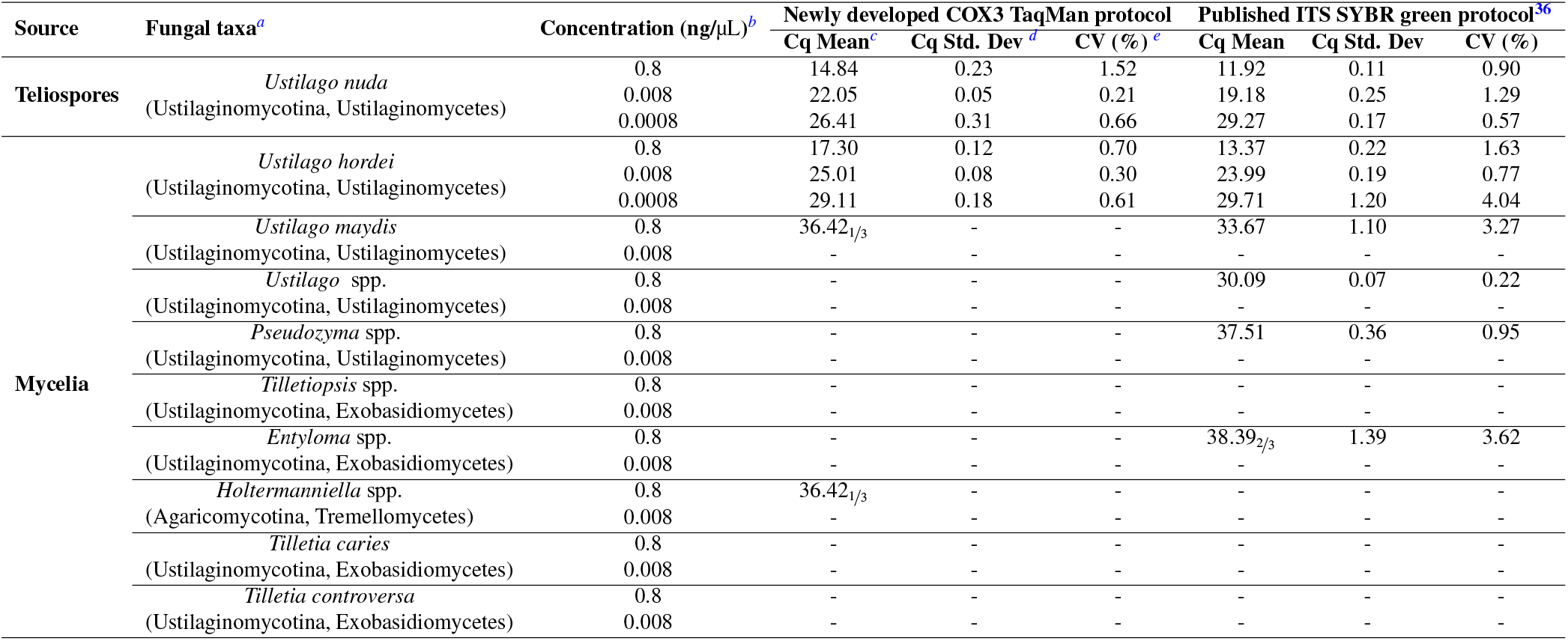
The assessment of COX3 and ITS primer specificity for *Ustilago nuda*. The newly designed COX3 primers in multiplex and the published ITS primers in singleplex were tested with DNA extracted from mycelia and teliospores of the target (*U. nuda*) and non-target fungal species (all other mycelial isolates listed that come from species other than *U. nuda*). Three technical replicates were used for both protocols. The number of amplified replicates is denoted as a subscript in the quantification cycle (Cq) mean when fewer than three amplified. The higher Cq value indicates less amplification. Samples with no amplification are indicated by the symbol “-”. ^*a*^Taxonomic classification with subphylum and class in parenthesis ^*b*^Final DNA concentration in each qPCR reaction ^*c*^Average cycle threshold of triplicates ^*d*^Standard deviation of the quantification cycle among triplicates ^*e*^Coefficient of variation (Cq Std. Dev.*/*Cq Mean)*100

**Figure 1.**
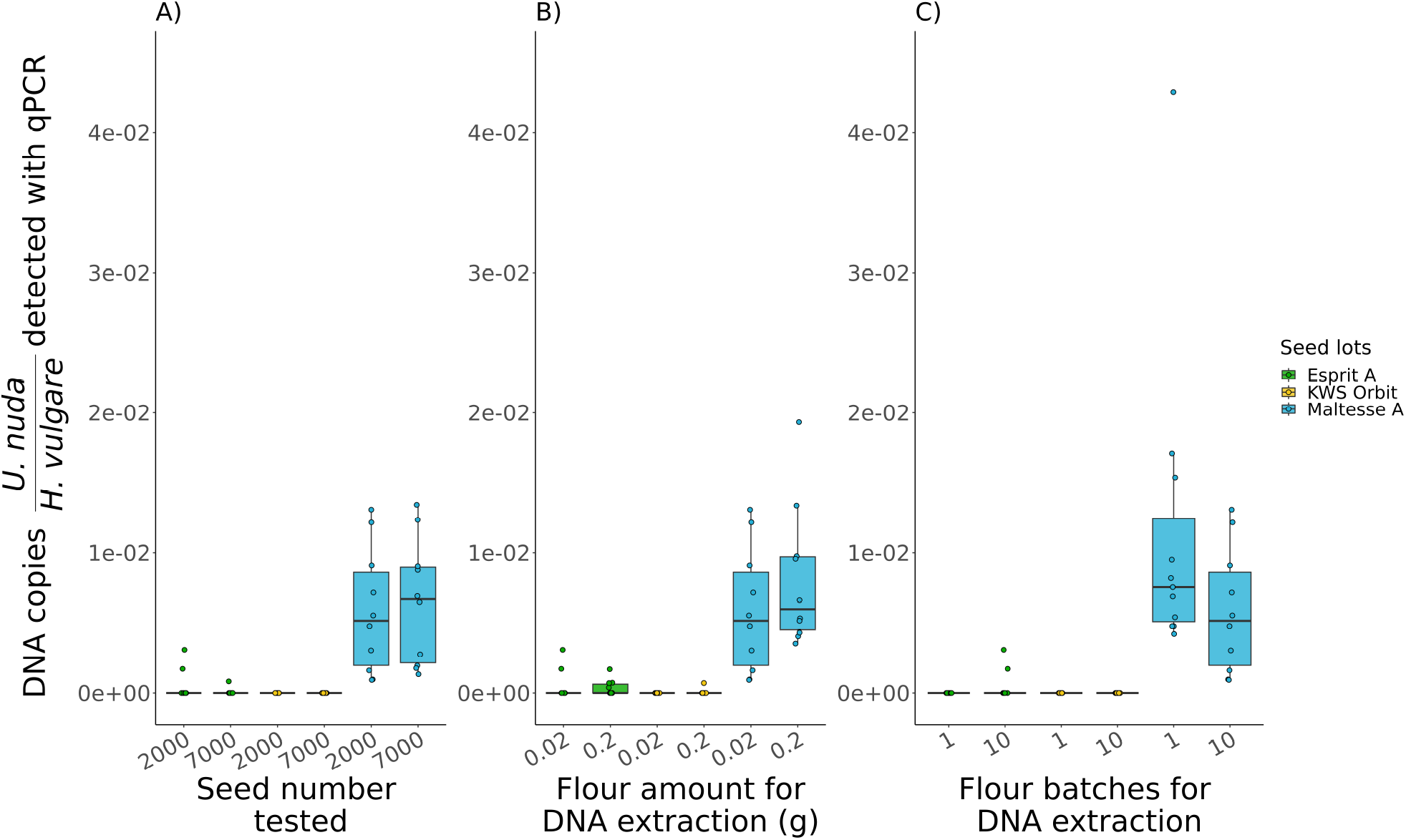
Key parameters evaluated to establish a representative *Hordeum vulgare* seed lot sample for *Ustilago nuda* qPCR detection. The parameters included: (A) the number of seeds tested (2000 versus 7000), (B) the amount of flour used for extraction (0.02 g versus 0.2 g), and (C) the number of flour batches used for the ten DNA sub-sample extractions (i.e. one batch extracted ten times or ten batches extracted one time each). The later analysis was used to assess inter- and intra-flour batch *U. nuda* DNA variation. The parameters in (A) and (B) were assessed with ten DNA extractions, each from a separate flour batch. *U. nuda* infection levels were quantified with the qPCR method developed in this study using the ratio *U. nuda* to *H. vulgare* DNA, which normalizes the pathogen DNA to its host DNA. Three different *H. vulgare* seed lots, each of a different cultivar, are shown in different colors. Kruskal-Wallis tests showed no significant differences in the detected *U. nuda* infection levels between each of the three key parameters within cultivars.

We tested non-target plant DNA with our newly developed GADPH primers to determine which plant species’ DNA would be suitable to include in the non-template DNA control and the standard curves. The DNA extracted from barley (*H. vulgare*), wheat (*Triticum aestivum*), and lentil (*Lens culinaris*) seeds were tested in our multiplex reaction. The primers successfully amplified barley DNA (Cq = 24.97*±* 0.24), but they showed reduced amplification of wheat DNA (Cq = 34.53 *±* 0.37), and no amplification of lentil DNA, demonstrating that lentil can be used as the non-template DNA in the barley standard curve and control. The data acquired for the main experiments in this paper showed minimal variation of *H. vulgare* detection based on the Cq values of the HEX-labeled *H. vulgare* GADPH probe (Supplementary Information, Tables 3, and 2)

**Table 2.**
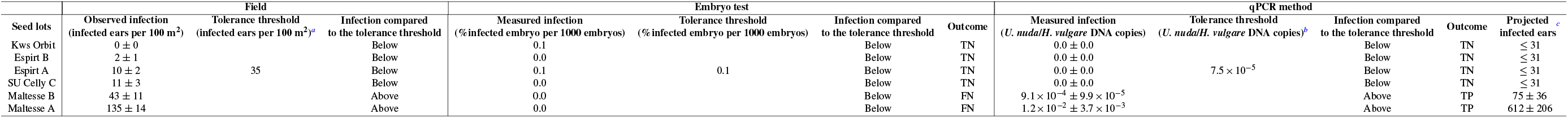
Comparison of *Ustilago nuda* infections observed in the field to those measured using the laboratory detection methods, the embryo test and qPCR method, on six commercially available seed lots. The field infection levels of each seed lot were evaluated in two years from four replicate plots per year, and these values served as a reference for the true infection levels. The percentage of infected embryos was based on one evaluation of 1000 seeds per seed lot. The infections detected with qPCR were based on ten extractions per seed lot, each with 0.02 g of flour per extraction. The flour for the ten extractions was obtained from 2000 milled seeds of the same seed lot. The standard errors are shown for the field observations and qPCR measured infection. All measured infection levels were determined to be either above or below the method-specific tolerance threshold for each seed lot, and these binary classifications were then compared with confusion matrices (Supplementary Information, Table 7). The binary classification for the field observations served as the actual values, and for the laboratory detection method, it served as the predicted values. The predicted values were then categorized as true positive (TP), true negative (TN), false positive (FP), or false negative (FN) depending on whether they reflected the actual values. These categorizations are reported in the outcome column. ^*a*^Tolerance threshold (infected ears per 100 m^2^) calculated from the number of infected plants derived from 0.1% infected seeds at a sowing density of 350 seeds/*m*^2^. ^*b*^Threshold (*U. nuda*/*H. vulgare* DNA copies) corresponding to 35 infected ears/100*m*^2^ and derived from the relationship *U. nuda* qPCR assay results with observed field infections (Figure2) ^*c*^Projected infected ears: estimated from the measured infection by qPCR and its corresponding field infection level according to their linear relationship (Figure2)

**Table 3.** Performance evaluations of the embryo test and our qPCR method in the detection of *Ustilago nuda* infections across six commercially available seed lots. The performance metrics include sensitivity, specificity, precision, false discovery rate, and accuracy. These performance metrics were based on the six seed lots’ agreement between their predicted values, which classified the seed lot as above or below the method-specific tolerance threshold, and their actual values based on the classification of the infections observed in the field as above or below its tolerance threshold (Table 2). The outcome of each seed lot’s classification according to the confusion matrices (Supplementary Information, Table 7) was used to evaluate the laboratory methods’ performance. Metrics marked with “-” were incalculable due to the absence of positive test results in the embryo test.

|  | Sensitivity | Specificity | Precision | False discovery rate | Accuracy |
| --- | --- | --- | --- | --- | --- |
| Embryo test | 0.0 | 1.0 | - | - | 0.6 |
| qPCR method | 1.0 | 1.0 | 1.0 | 0.0 | 1.0 |

### A representative seed lot sample for *Ustilago nuda* qPCR detection

Our qPCR method relies on DNA samples extracted from milled seeds, so we evaluated different sample preparation procedures to ensure an accurate seed lot representation. We tested three key parameters to establish a representative seed lot sample: 1) the number of milled seeds, 2) the amount of seed flour for DNA extraction, and 3) the number of milled seed batches. The evaluation of these key parameters was conducted on three seed lots, each from a different cultivar: Esprit, KWS Orbit, and Maltesse. The qPCR results, based on the ratio *U. nuda/H. vulgare* DNA copies, showed no significant differences between any of the parameters tested within each cultivar (Figure 1, Supplementary Information, Table 4). However, wide variability in infection levels was observed among subsamples, especially as the average infection level of the seed lot increased (Supplementary Information Table 4).

**Table 4.**
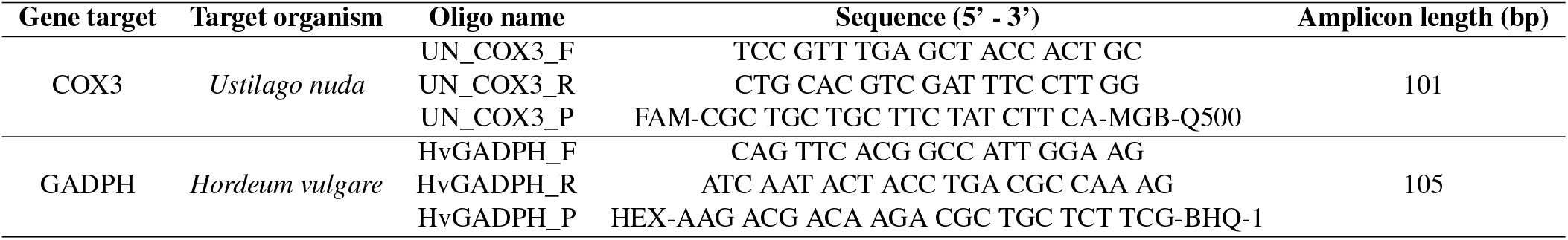
Primers and probes used in this study. The primers and probes target the plant pathogen, *Ustilago nuda*, and its host, *Hordeum vulgare*. The COX3 probe has a minor groove binder (MGB) modification on its 5’ end.

### The relationship between laboratory detection methods and field results

The relationships between *U. nuda* infection levels observed in the field and those measured by laboratory detection methods, including the embryo test and the qPCR method, were evaluated with three barley cultivars: Azrah, Semper, and SU Celly. These seed lots were chosen for this experiment based on our access to naturally infected seeds with a high infection level. Samples of each cultivar were composed of four different infection levels, which were achieved by using naturally low-infected (certified seed lots) and high-infected seed lots (based on observed field infection levels of the previous generation) and mixing them. The two mixtures for each cultivar were made with either 10% or 1% from the high-infected seed lot and 90% and 99% from the low-infected lot, respectively, totaling twelve seed samples. The two naturally infected seed lots and their mixtures with different infection levels were used to study the correlation between field observations and laboratory detection methods. Additionally, this relationship was used to derive a tolerance threshold for the qPCR method. To determine which model type best fits the relationship between the qPCR measurements and the field observations, we tested three models (linear, polynomial and exponential) and compared them (Supplementary Information, Table 5). The linear model was chosen for its predictive performance (highest adjusted *R*^2^) and lower complexity (lowest AIC and BIC) compared to the polynomial and exponential models. The positive linear relationships showed that the laboratory detection methods are able to predict the field infection level (Figure 2) and that the correlation is about 22% higher in the qPCR method compared to the embryo test based on the adjusted *R*^2^ values. The correlation between the qPCR measurement and the field observations was then used to develop a tolerance threshold for the qPCR method. The tolerance threshold used in our study was 35 infected ears based on 350 seed/*m*^2^ sowing density (equivalent to 0.1% infected embryo and the assumption that one infected embryo develops one infected ear, Supplementary Information, Figure 2). This tolerance threshold correlated with 7.50 *×* 10^*−*5^ *U.nuda/H. vulgare* DNA copies.

**Figure 2.**
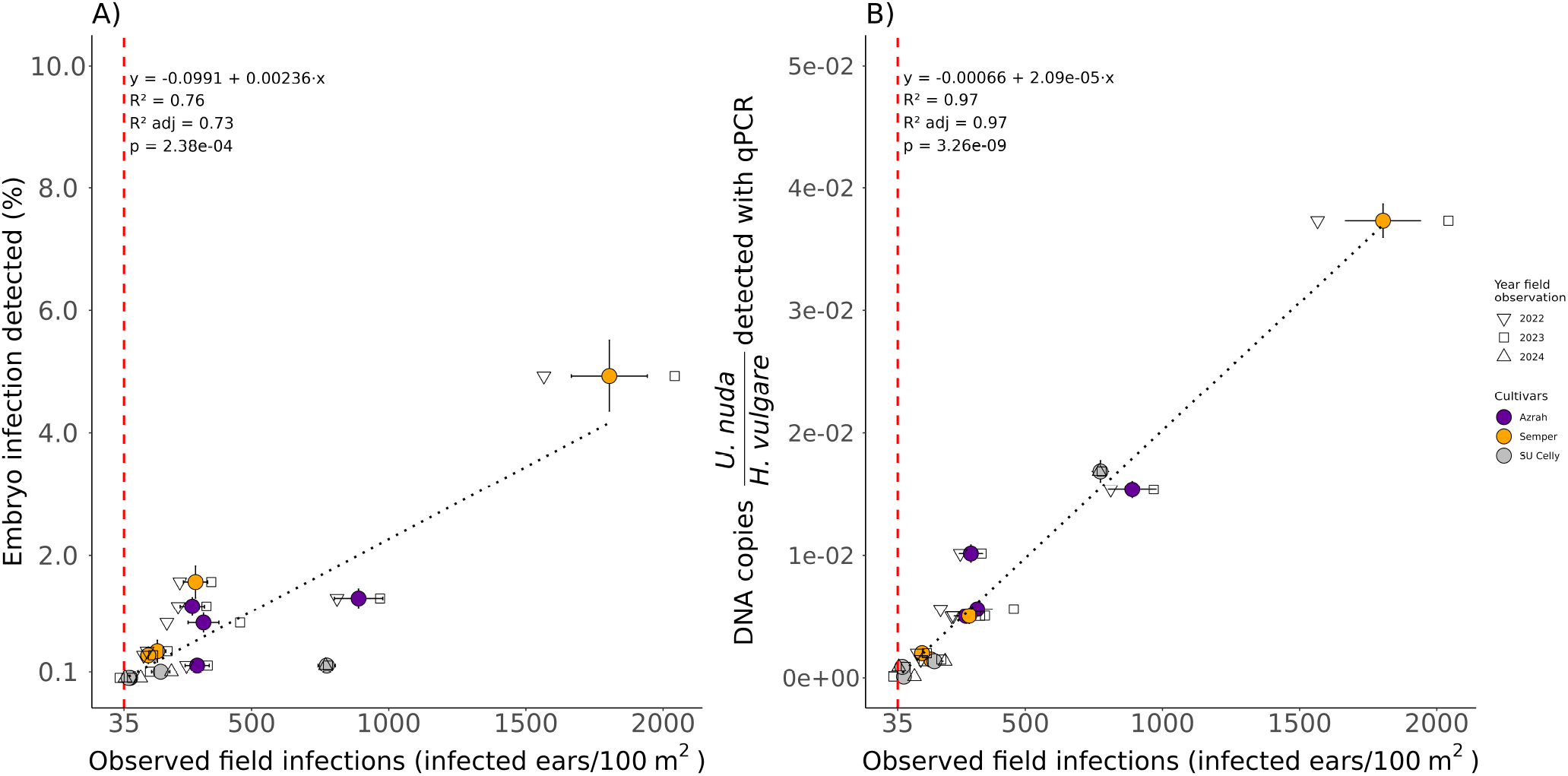
Relationship between the *Ustilago nuda* infection observed in the field and measured with laboratory detection methods: (A) embryo test and (B) qPCR method. Observed field infections, used as the reference infection level for each sample obtained from naturally infected seed lot or their mixtures, are plotted on the x-axis as the independent variable. The *U. nuda* measurements from the laboratory detection methods that are evaluated in this study are plotted on the y-axis as dependent variables. The linear models’ lines of best fit between field observation and the laboratory detection methods are illustrated with black dotted lines. The linear model equations, *R*^2^, adjusted *R*^2^ (*R*^2^ adj), and p-value are shown in the top left corner of each plot. Each sample from each cultivar was sown in two of the three field growing seasons. The different shapes indicate the two years of observed field infections for each sample, and the average infections of each sample are shown as a circle. In both plots, the red vertical dashed line indicates the field tolerance threshold value (35 ears per 100 *m*^2^).

### The comparative performance of laboratory *Ustilago nuda* detection methods to predict disease levels in the field

The qPCR method and the embryo test were evaluated for their ability to detect *Ustilago nuda* infections in six commercially available *Hordeum vulgare* seed lots from four cultivars: Esprit (two seed lots, A and B), KWS Orbit, Maltesse (two seed lots, A and B), and SU Celly (seed lot C). Three of the seed lots —Esprit A, KWS Orbit, and Maltesse A —were the same used to establish a representative seed lot sample for the *U. nuda* qPCR detection method. Each seed lot was sown in four 20 *m*^2^ field plots in two growing seasons. The observed field infection levels across the two growing seasons were averaged and served as the reference for the performance of the embryo test and qPCR method to detect *U. nuda* in seed (Table 2). The field infection levels ranged from 0.00% to 0.44% across all of the seed lots in both growing seasons (Supplementary Information, Figure 2). We created confusion matrices to compare the performance of the embryo test and qPCR method. For each matrix, we classified the infection levels measured by the embryo test and qPCR method as either above or below their respective tolerance thresholds: 0.1% infected embryos and 7.50*×*10^*−*5^ *U. nuda/H. vulgare* DNA copies. These classifications served as the predicted values (Supplementary Information, Table 7). In parallel, the seed lots’ infection levels were classified as above or below the tolerance threshold based on their observed infected ears in the field (Table 2), and these were used as the actual (or true) values. Based on the comparisons between the actual values and predicted values, we assessed which samples they accurately categorized to have infection levels above or below their tolerance thresholds (Table 2, Supplementary Information, Table 7). Based on the infection detected by the embryo test, all tested seed lots were classified as having infection levels below the embryo test’s tolerance threshold (0.1%). However, two of these seed lots had field infection levels exceeding the field’s tolerance threshold, so the embryo test produced false negatives by failing to detect those infections. The seed lot infections detected using the qPCR method, on the other hand, were correctly identified as above or below its tolerance threshold and matched the actual values.

We then used the confusion matrices to evaluate the sensitivity, specificity, precision, false discovery rate, and accuracy of the *U. nuda* detection methods in six commercially available seed lots. These attributes served as performance metrics. Together, the performance metrics showed that the qPCR method classified the *U. nuda* infection levels better than the embryo test (Table 3). Two of the six seed lots had infection levels exceeding the field’s tolerance threshold so have actual values that are above its threshold (Table 2). The qPCR method showed a sensitivity and specificity of 1.0, accurately identifying the infection of these two seed lot infections as above its tolerance threshold and the other four seed lots as below it. On the other hand, the embryo test identified the infection of these two seed lot samples as below its tolerance threshold when they should be above it. Because the embryo test yielded no predicted values that were above its tolerance threshold, we cannot calculate the embryo test’s precision and false discovery rate in this study. However, the embryo test successfully identified the seed lots with actual infection values below the field tolerance threshold, resulting in a specificity of 1.0 and a sensitivity of 0.0. Overall, the qPCR method consistently detected *U. nuda* infection levels more accurately and precisely in the evaluated six commercially available seed lots than the embryo test (Table 3).

## Discussion

Seed health tests help prevent seedborne disease outbreaks and reduce reliance on plant protection products (PPPs)^37^ because they indicate whether untreated seed lots present low risk of disease outbreaks. However, traditional methods —such as field inspections and visual seed assessments —are often time-consuming, resource-intensive, and prone to inaccuracies^25,26,38^. These challenges are particularly problematic for detecting cryptic seedborne infections, such as *Ustilago nuda* in barley (*Hordeum vulgare*), in which infected seeds are asymptomatic^39^, making it difficult to determine whether seed lots require treatments or should not be sown. To address these limitations, we developed a multiplex qPCR method that quantifies *U. nuda* DNA relative to host DNA in bulked milled seed samples, thereby improving both detection accuracy and scalability.

We tested barley seed lots with our qPCR method and the existing validated test, the embryo test, to evaluate their ability to detect *U. nuda*. The direct comparison of a new qPCR method to an existing test can be problematic; if the validated test performs poorly, it provides an unreliable indicator of the seed lot’s infection status. Because of the drawbacks associated with directly comparing detection methods, we used the observed number of infected ears in the field as a reference to determine the true *U. nuda* infection rate in seed lots. Based on the adjusted *R*^2^ values in our study, the embryo test explained 24% less variation in the field infection rates compared to our qPCR method. The difficulty of visualizing *U. nuda* mycelia with the embryo test may have contributed to its reduced ability to predict field infection levels. Therefore, the seed inspectors may underestimate the seed infection levels (Supplementary Information, Table 8). Although recent protocol improvements to the embryo test, including methyl blue staining, have been introduced to enhance mycelium visibility^24^, these modifications were not tested in our study. However, these protocol adjustments still do not overcome the longer working time required to obtain results compared to a DNA-based test.

Molecular based detection methods on seed can offer higher sensitivity and specificity with faster throughput compared to visual seed inspections^27,28^. Our multiplex qPCR amplifies both the host and the pathogen DNA to improve the relative quantification of *U. nuda* detection in seed. Previous studies have amplified host DNA as an external control in separate qPCR reaction or within a multiplex reaction as an internal control to confirm the quality of the DNA extraction and PCR amplification^31,32^. However, their protocols have not used host DNA to normalize the pathogen DNA. Our qPCR method not only uses an internal control of host DNA, but it also provides a means to normalize the pathogen DNA. The normalization step is particularly useful to quantify an internal embryo infection, such as *U. nuda*, that is asymptomatic and lays dormant in the seed as mycelia. The normalized *U. nuda* DNA levels to *H. vulgare* DNA in each sample enhances the reliability and robustness of *U. nuda* detection because it reduces the reliance on quantification cycle (Cq) values, also known as threshold cycle (Ct) values. The use of Cq values as a cutoff for pathogens’ detection does not account for possible variability in DNA extraction efficiencies and intra- and inter-plate differences^40,41^.

The normalization procedure with the host’s DNA is particularly relevant for *U. nuda* because its mycelial biomass in the embryo, can vary among seeds, and this variability is not considered during the visual embryo test. Moreover, most seedborne pathogenic fungi are quantified as spores or colonies per seed without destructive sampling methods^29,31,42^, which is not possible for *U. nuda* due to its location in the seed embryo. Therefore, when pathogenic fungi are measured as spores or colonies per seed based on visual assessments, it is easier to assess the infection rate of the seed and observe the subsequent field infection levels. However, due to the *U. nuda* location within the seed, this relationship between seed and field infection levels is difficult to surmise. Additionally, the tolerance level of an *U. nuda* infection in seed is low —for example, 1 in 1000 is the tolerance threshold for many countries^20,22^ —so that destructively sampling enough seeds to estimate the field infection rate quickly becomes labor-intensive and cumbersome. Moreover, the relationship between the *U. nuda* mycelial biomass and the number of infected ears in the field remains unknown because each tested seed requires its destruction. To address the challenge to related seed and field infection levels, we established equivalent tolerance thresholds for the field, the qPCR method and the embryo test based on the assumption that one infected seed produces one infected ear. This assumption is also supported by our field observations (Supplementary Information, Figure 2B).

We used a tolerance threshold to assess the ability of our qPCR method and the embryo test for predicting field infections in commercially available seed lots. Although both laboratory detection methods showed high specificity, the qPCR method was more sensitive because it detected infections that the embryo test missed. These findings suggest that the embryo test may underestimate *U. nuda* infections, potentially failing to prevent its spread and future economic losses. Additionally, the qPCR results closely matched the observed infections in the field. One of the tested seed lots had an infection level close to the tolerance threshold, and it showed concordance between the observed field infections and the measured *U. nuda* infection with qPCR: both measurements were not consistently above or below their respective tolerance threshold (Supplementary Information, seed lot Maltesse B in Tables 6 and 9).

Overall, our qPCR method showed a higher precision and accuracy than the embryo test. The embryo test’s lower performance can be attributed to its detection limit of 0.1% infected embryo based on analyzing 1000 embryos per sample. However, the International Seed Testing Association suggests increasing the sample size to 2000 – 4000 embryos to decrease the test’s detection limit to 0.05% —0.025%^24^. Doubling or quadrupling the sample size would add time and cost, making the test impractical for large-scale applications^25^. In contrast, we can predict down to 31 infected ears per 100 *m*^2^ at a sowing density of 350 seeds/*m*^2^ with our qPCR method, which corresponds to an embryo infection rate of 0.009%. We reached this improved detection sensitivity because our qPCR method uses pooled seed samples to enable the detection of even small amounts of *U. nuda* DNA. Indeed, qPCR-based detection methods for other seedborne pathogens show the advantages to pooling seed samples^28–32,43^. Additionally, unlike previous *U. nuda* molecular detection methods that involve single embryo extraction or labor-intensive techniques (e.g., mortar and pestle)^10,33–35^, our approach involved seed milling to pool seed samples. This sample processing facilitates potential large-scale applications and improves detection at low infection rates^32,34,35,44^.

Despite the potential for high-throughput, we noticed that our qPCR method overestimated the projected number of infected ears in seed lots that were above the field tolerance threshold (Table 2). This overestimation may be reduced if more seed lots are tested for the correlation between qPCR results and field observations (Figure 2). Another possible explanation is that highly infected seed lots may contain infected seeds with more mycelial biomass, which could increase the number of infected ears per plant (Supplementary Information, Figures 2). To eliminate the possibility of non-specific amplification contributing to the overestimation of infected, we investigated the species-specificity of our qPCR method. Our qPCR method showed improved specificity compared to the most recently published qPCR protocol to detect *U. nuda*^36^. We consistently amplified *Ustilago hordei* (covered smut of barley) with our qPCR method, but no other fungal species isolated from Swiss seed lots. Since *U. hordei* and *U. nuda* cause similar damage and the field tolerance threshold is based on the number of infected ears of either pathogen in the field, the *U. hordei* amplification does not compromise the practical application of our qPCR method^21^. Another common limitation in qPCR methods is the presence of inhibitors that can cause false negatives^45^. Although we did not include any additional inhibitor-removal steps during DNA extraction^32^, our tests showed no inhibition in detecting *U. nuda* in seed (Supplementary Information, Table 10). To help validate the method’s reliability and facilitate its adoption in routine seed health testing, ring testing across multiple laboratories and sowing seed lots under diverse conditions would further ensure the robustness of our qPCR method^46–48^.

In conclusion, our qPCR method for untreated barley seed provides a more accurate, scalable, and practical alternative to the existing validated seed health test for detecting *U. nuda* in embryos. It serves as a robust secondary check on harvested seed certified through field inspections; rather than relying on visually infected ears in field, which do not necessarily capture the next generation’s infection rate, it detects the actual infection in harvested seed. The combination of field inspections on plants grown from first-generation certified seed and our qPCR method on second-generation seed will help farmers and seed producers minimize the *U. nuda* infection in their barley fields. Additionally, rather than the use of prophylactic seed treatments, the incorporation of more accurate seed health testing prior to treatments can improve targeted PPP application. Targeted seed treatments follow the principles of integrated pest management and support disease management practices within the seed sector^1^.

## Methods

### Primer and probe development for multiplex qPCR to detect *Ustilago nuda*

Primers and probes for the multiplex qPCR reaction were developed to specifically target the cytochrome c oxidase subunit III (COX3) gene from the plant pathogen *Ustilago nuda* (GenBank accession number: HQ013017) and the glyceraldehyde-3-phosphate dehydrogenase (GAPDH) gene from its host, *Hordeum vulgare* (GenBank accession number: MT933276). The primers were designed using the Primer3 software tool^49^ as a basis for primer selection (Table 4). The potential amplification of non-*U. nuda* species was first checked using the NCBI nucleotide database and Primer-BLAST^50^. All primers and probes were synthesized by Microsynth (Balgach, Switzerland). The annealing temperatures for the primer and probe sets were first optimized in the range of 58 ^*°*^C to 68 ^*°*^C in singleplex reactions, followed by a test gradient of 63 ^*°*^C to 66 ^*°*^C for multiplex reactions. Both singleplex and multiplex reactions demonstrated comparable amplification efficiencies: 99% for COX3 and 90% for GAPDH.

The newly developed *U. nuda* primer sets were compared to the most recently published *U. nuda* primers^36^, which amplifies the ITS region of *U. nuda* in a singleplex reaction using SYBR Green dye. The published ITS protocol was adapted for the Opus qPCR system (BioRad, USA) with a reaction volume of 10 µL, and reactions were run using the GoTaq qPCR Master Mix (Promega, USA). To evaluate the specificity of the *U. nuda* primers, both protocols were tested on DNA extracted from mycelia of various basidiomycetous fungal species (Table 1). The qPCR protocols with either our newly developed COX3 or the published ITS primers were run using a range of DNA concentrations, including 0.8, 0.008, and 0.0008 ng/µL. The positive control for *U. nuda* originated from teliospores collected from the *H. vulgare* cultivar Cassia during a 2022 field trial at Reckenholz (ZH), Switzerland.

An *in silico* analysis with the COX3 primers indicated the highest likelihood of cross-amplification with *Ustilago hordei* (Supplementary Information, Figure 1). To check potential cross-amplification, *U. hordei* mycelia (strain Mat1, provided by Gunther Doehlemann, University of Cologne) was tested in qPCR reactions with the same DNA concentrations as *U. nuda*, 0.8, 0.008, and 0.0008 ng/µL. DNA from non-target basidiomycetous mycelia was tested at concentrations of 0.8 and 0.008 ng/µL in qPCR reactions. Non-target basidiomycetes —*Ustilago maydis, Tilletia caries, Tilletia controvera, Tilletiopsis* spp., *Pseudozyma* spp., another *Ustilago* spp. with teliospores approximately 9 *µ*m in diameter, *Entyloma* spp., and *Holtermanniella* spp. —were included in this analysis to assess primer specificity. *Tilletiopsis* spp., *Pseudozyma* spp., another *Ustilago* spp., *Entyloma* spp., and *Holtermanniella* spp. were isolated from Swiss seeds or smutted ears (Supplementary Information, Subsection Supplementary Methods), and their identity was deduced based on sequence similarity to other Basidomycetes using BLAST searches against the core nucleotide database and whole genome shotgut contigs (Supplementary Information, Table 11). DNA from *T. caries* and *T. controversa* was extracted from single spore mycelial isolates from bunt-infected seeds collected during a 2022 pot trial in Reckenholz (Zurich, Switzerland) and a 2023 field trial in Unterwasser (St. Gallen, Switzerland), respectively. The *U. maydis* strain FB2 was provided by G. Doehlemann. Sanger sequencing of the ITS1 region confirmed the species identity of the fungi isolated in this study (NCBI Accession numbers to be provided upon publication). DNA extractions of the mycelia and teliospores were performed using the NucleoSpin Plant II Mini kit (MACHEREY-NAGEL, Germany), and DNA concentration was quantified with Nanodrop (Thermo Fisher Scientific Inc, USA). Primers and probe targeting the *H. vulgare* GAPDH gene were tested to determine whether the plant material DNA from wheat (*Triticum aestivum*) or lentil (*Lens culinaris*) seed could be used as the control and the standard curve’s non-template DNA.

### Seed material and subsample preparation

Untreated naturally infected seeds from two-row and six-row barley cultivars were used in this study. For the experiment to test a representative seed lot sample, we used three commercially available seed lots from the cultivars: Esprit, Maltesse, and KWS Orbit. To assess the correlations between the observed field infection level and *Ustilago nuda* detected in seeds with the embryo test and qPCR method, we focused on three different cultivars: Azrah, Semper and SU Celly. For each of these cultivars, we used two seed lots including a more infected and less infected lot. The more infected lots were mixed with the less infected lots in the following proportions: 10:90 and 1:99. The original two seed lots plus these two mixtures resulted in four infection levels for each cultivar, which were then sown in the field and analyzed using the qPCR method and the embryo test. Next, we conducted an experiment to identify whether the results from the qPCR method or the embryo test on barley seed provided a more accurate prediction of *U. nuda* infection levels in the field. For this experiment, we used the same three commercially available lots used to establish a representative seed lot sample and three additional commercially available seed lots from the cultivars, Maltesse, Esprit, and SU Celly. The commercially available SU Celly seed lot used in this part of the study differed from the two seed lots used in the previous experiment explained above.

In all experiments, the seeds from each 25 kg seed lot were placed in 12 cm x 60 cm x 80 cm containers so that the subsamples could be collected randomly. Sufficient subsample quantities for all field trials and laboratory detection methods (i.e. embryo test and qPCR method) were collected together at the beginning of our study to ensure that the same subsample was used throughout the study. For each seed lot and seed lot mixtures, 200 g of seed (approximately 4000 seeds) were sent to the Mycology Laboratory at the Bavarian State Research Center for Agriculture, where the embryo test was performed. For the qPCR method, the seeds —2000 or 7000 seeds depending on the test conducted —were milled (CT 293 Cyclotec; Labtec Line, Denmark). To avoid cross-contamination a thorough cleaning of the mill with compressed air was performed between each seed sample.

### Field trial set-up

The field trials were conducted at Agroscope in Zurich, Switzerland over two growing seasons for each seed lot. The trials took place during 2021-2022 and 2022-2023 for the cultivars Azrah and Semper, while SU Celly and the six commercially available seed lots were sown for the growing seasons 2022-2023 and 2023-2024. Each growing season, the field location was changed to comply with crop rotation practices. Seeds were sown in the field during the first week of October (Supplementary Information, Figure 3). Each seed lot and seed lot mixture was sown with four replicates in completely randomized blocks. Seeds were sown at a density of 350 seeds/*m*^2^ and at a depth of 3-4 cm.

Seed lots and their mixtures, used to establish the correlation between the results of the laboratory detection methods and the observed field infection levels, were sown in 9.0 *m*^2^ plots. Prior to the flowering stage, approximately 0.65 m from the short ends of each plot was trimmed with a motor mower. The shortened plots facilitated the assessment of infections and ensured plot separation. Following the trimming, the average final plot size was 5.2 *m*^2^. Commercial seed lots, used to evaluate the laboratory methods’ performance for *Ustilago nuda* detection, were sown in larger plots of approximately 20 *m*^2^. Gaps were left between plots during the sowing to ensure consistency in plot size. Plot dimensions were measured at plant maturity, and the average plot size was 19.4 *m*^2^.

At the second leaf stage (Supplementary Information, Figure 3), germination rates were measured by counting seedlings along a linear meter multiple times within each plot. To minimize boundary effects, the counts were taken at two points along the second and sixth rows (of seven rows) and approximately 1 meter from the short plot edges. In the larger plots, this procedure was slightly adjusted, and counts were conducted at six points per plot instead of two to account for the size difference. During the flowering stage, ears were counted in a meter long frame that covered an area of 0.3 *m*^2^. For the ear counts, plot edges were avoided, and ears were counted in the second and sixth rows (of seven rows). Two and six sampling points per plot were used for the 5.2 *m*^2^ and 19.4 *m*^2^, respectively. From this ear count, the total number of ears per plot was estimated by averaging the number of ears within the each 0.3 *m*^2^ sampling area and scaling it to the measured plot size. In the flowering stage (Supplementary Information, Figure 3), the total number of infected ears was recorded within each entire plot. The number of infected ears was divided by the estimated total number of ears to calculate each plot’s infection rate.

### Embryo test implementation

The International Seed Testing Assosiation (ISTA) method “Detection of *Ustilago nuda* in *Hordeum vulgare* subsp. *vulgare* (barley) seed by dehulling and embryo extraction”^24^ was used for the embryo test. This examination was conducted on 1000 embryos from the seed lots and seed lot mixtures that had been sown in the field and tested with the qPCR method. For the seed samples of the cultivars Azrah and Semper, the same extracted embryos were examined by three different evaluators (Supplementary Information, Table 8).

### Sample preparation for the DNA extraction

To establish a representative flour sample from each seed lot, a series of tests was conducted on the following milled commercial seed lots: Maltesse A, Esprit A, and KWS Orbit. The optimal seed quantity for milling and the appropriate amount of flour for DNA extraction were assessed using ten subsamples each collected from ten different milled flour batches. Additionally, the uniformity of infection across milled flour batches was evaluated by extracting ten subsamples from either one or ten milled flour batches. For the seed quantity test, we analyzed two sample sizes: 2000 seeds and 7000 seeds. The 2000 seed sample corresponds to the number of seeds used in the working sample of the embryo test^24^, while the 7000 seed sample corresponds to the number of seeds sown in a field trial in 20 *m*^2^ plot and sowing density of 350 seed/*m*^2^. To establish the flour amounts required for DNA extraction, 0.2 g and 0.02 g samples of 2000 milled seeds were used; the number of seeds was selected based on the previous test’s results that showed no significant difference between 2000 and 7000 seeds in *Ustilago nuda* detection. To assess the uniformity of infection within and between flour batches, DNA was extracted from 0.02 g of flour samples obtained by milling 2000 seeds; the amount of flour was established in the previous test that showed no significant difference in *U. nuda* detection between the two flour amounts. These flour samples were collected in two ways: ten times from a single flour batch, and once from ten separately milled flour batches. Based on this series of tests, a sampling protocol was established that consisted of milling 2000 seeds and extracting DNA from ten 0.02 g subsamples of flour from a single batch.

### Seed sample DNA extraction

The milled seed samples of 0.02 g and 0.2 g were extracted using the NucleoSpin 96 Plant II Mini kit (MACHEREY-NAGEL, Germany) according to the manufacturer’s protocol with the following modifications. For both flour amounts, a 3 mm tungsten bead (Retsch, Germany) was added to facilitate mechanical homogenization during the extraction process. For the extractions of 0.02 g flour, the flour was added to the NucleoSpin lysis tube strips. The initial homogenization and mechanical destruction of the sample was performed with the Tissue Lyser II (Qiagen, Germany) at a rate of 30 1/s for four consecutive 1-minute intervals, and following each interval, the orientation and position of the samples were changed. Cell lysis was performed with 600 µL of PL1 buffer provided with the kit, and the samples were mixed four times with the Tissue Lyser II (Qiagen, Germany) before adding 12 µL of RNase A (concentration 12 ng/µL). Subsequently, the samples were gently mixed and placed for 1 hour at 65 ^*°*^C in a LAUDA E200 heated water bath (Lauda Dr. R. Wobser GMBH & CO. KG, Germany) and then gently mixed. Following the washing steps as described in the manufacturer’s protocol, each sample was eluted two times with 100 µL of elution buffer from the kit, yielding a final volume of 200 µL.

For the extractions of 0.2 g flour, further modifications were taken in the cell lysis step to account for the increased volume of material. A 50 ml Falcon tube was used for the homogenization and mechanical disruption step, which was conducted with the Bead Ruptor 24 (Omni International, USA) at a speed of 2.1 m/s for three 1-minute intervals. After each 1-minute interval, the sample orientation and position were changed. The amounts of PL1 buffer and RNAse A were increased to 4.03 ml and 66.78 µL, respectively, to account for the increased sample dry weight. The three different sized NucleoSpin Plant II kits’ maximum recommended sample dry weights were used to interpolate the reagent volumes for the 0.2 g sample. Following centrifugation, 400 µL of lysate was transferred to the next step according to the kit’s instructions for extractions of 0.02 g of plant material. The remainder of the extraction was carried out as described as above for the 0.02 g flour DNA extraction.

### *Ustilago nuda* and *Hordeum vulgare* DNA quantification

A series of gBlock double-stranded synthetic oligonucleotides fragments (Integrated DNA Technologies, USA) was prepared with a 5-fold serial dilution to create a standard curve for absolute quantification (Supplementary Information, Table 12). Distinct 125 bp gBlock fragments were synthesized for both the *Hordeum vulgare* and the *Ustilago nuda* targets. These fragments included 12 bp and 10 bp flanking regions on both sides of the COX3 and GAPDH amplicons, respectively. The gBlock fragments were initially diluted to a concentration of 0.03 ng/µL and stored at −20 °C with 0.2 mg/mL Poly(A) (Sigma-Aldrich, Germany). The diluted gBlock fragments were then divided into aliquots suitable for a single plate to limit freeze-thaw cycles. The standard curve copy numbers, calculated from the molecular weight of the gBlock fragments, ranged from 62500 to 0.8 for COX3 and from 1563000 to 500 for GAPDH.

To replicate the PCR reaction conditions, template DNA from uninfected *H. vulgare* seedlings was included in the *U. nuda* standard curves, and *Lens culinaris* flour was added to the *H. vulgare* standard curve. The final amount of DNA in the standard curves was 50 ng in each reaction. Each qPCR plate included technical triplicates of the standard curves, experimental samples, and controls. The controls run on each plate included water and the non-target template DNA (i.e. uninfected *H. vulgare* seedlings and *L. culinaris* seed for the COX3 and GADPH standard curves, respectively) that had been used in the standard curves.

DNA sample concentrations were measured using a Varian Cary Eclipse Spectrophotometer (Agilent, USA). Samples were diluted to a concentration of 12 ng/*µ*L using a Pipetmax Gilson 268 system in combination with Trilution software (Gilson, USA). The qPCR reactions were conducted in a final volume of 10 µL per well, containing 50 ng of DNA, 0.2 *µ*M of each COX3 primer and probe, 0.1 *µ*M of each GAPDH primer and probe, and 5 *µ*L of GoTaq Probe qPCR Master Mix (Promega Corporation, USA). The qPCR was performed using white 384-well plates sealed with adhesive optical sealing foil (Nolato TreffLab, Sweden). The thermal cycling conditions were initiated with a denaturation step at 95 ^*°*^C for 2 minutes, followed by 40 cycles of 95 ^*°*^C for 15 seconds and 65.3 ^*°*^C for 30 seconds. All reactions were run on the CFX Opus 384 Real-Time PCR System (Bio-Rad, USA). The gene copy numbers for COX3 and GAPDH in each sample were determined using CFX Maestro 2.2 software (version 5.2.008.0222) by comparing the quantification cycle (Cq) values to the corresponding standard curve.

The limit of detection (LOD) for *U. nuda* was established by running eight series of standard curve dilutions. Experimental samples were repeated when the standard deviation of the Cq values across triplicates exceeded 0.5 for either gene’s quantification. However, samples were not repeated if the average triplicate measurement for *U. nuda* was below the LOD for COX3, as the high standard deviation was likely due to the low levels of *U. nuda* target DNA. In cases where *U. nuda*’s COX3 measurement was below the LOD, *U. nuda* was considered to be undetectable, and a copy number of 0 was used in the statistical analysis. We chose to use 0 as the copy number for samples with undetectable levels of *U. nuda* to ensure consistency among the different detection methods (embryo test, qPCR method, and field observations). Both the field and embryo tests consider detection from 0, meaning that an undetectable result does not necessarily indicate the absence of infection, but rather an infection load below the detection threshold. The same baseline was used to provide a meaningful comparison to the reference (field) and embryo tests, which both start at 0. The infection level in each sample was calculated by taking the ratio of the mean copy number of *U. nuda* COX3 to the mean copy number of *H. vulgare* GAPDH.

## Data and statistical analyses

### Establishment of a representative seed lot sample for Ustilago nuda qPCR detection

All statistical analyses were conducted using R version 4.3.3. Three key sampling parameters were compared and analyzed with Kruskal-Wallis test to establish the sampling procedure used in the rest of the study. The tested parameters were: the seed number (2000 versus 7000), the flour amount (0.2 g versus 0.02 g), and the number of flour batches used for the ten DNA sub-sample extractions (i.e. one batch extracted ten times or ten batches extracted one time each).

### Determination of a tolerance threshold

The minimum number of embryos examined for the embryo test is 1000 per sample^24^. With a sample size of 1000 embryos, infections levels under 0.1% cannot be detected^25^. Based on the lowest infection rate possibly detected with the embryo test, a minimum field tolerance threshold was derived using the equation below. Although one embryo potentially results in zero or multiple infected ears, we used the most conservative number of infected ears produced by one infected seed. Therefore, we assumed that one infected seed produces one infected ear to calculate the field tolerance threshold. This assumption is also reflected in our observations (Supplementary Information, Figure 2). The expected infection level in the field, which was used as its tolerance threshold, was calculated from the minimum detectable infected embryos. To determine the number of infected ears per 100 *m*^2^ expected from an embryo test result of 0.1%, infected embryos, we used the equation below:

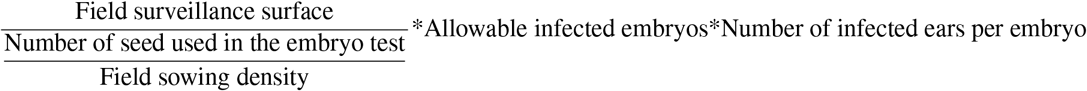

Based on our sowing rate of 350 seed/*m*^2^ and the use of the minimum detectable infected embryos, the field tolerance threshold was calculated to be 35 infected ears per 100 *m*^2^. The germination of all seeds was assumed because the embryo test does not account for a potential reduction in germination. This field tolerance threshold was used to derive the qPCR method tolerance threshold. To connect field infection levels to expected results of the qPCR method, we used the relationship between the naturally infected Azrah, Semper and SU Celly seed lots and their mixtures’ field data to their qPCR results. We used these data to derive a qPCR tolerance threshold that corresponds to 35 infected ears per 100 *m*^2^. Linear, polynomial and exponential correlations were tested to evaluate the relationship between the field observations and the qPCR method. The linear correlation was selected based on the adjusted *R*^2^, the mean squared error (MSE), the Akaike information criterion (AIC) and the Bayesian information criterion (BIC). The conversions of the tolerance thresholds from embryo test to the field observations and from the field observations to the qPCR method allowed us to indirectly compare both laboratory detection methods to each other.

### Performance evaluation of the laboratory detection methods: qPCR method and embryo test

The observed infected ears in the field were used as a reference to evaluate the performance of the laboratory detection methods, including the qPCR method and embryo test, to detect Ustilago nuda in six commercially available seed lots. We classified whether a seed lot was below or above each respective tolerance threshold: 0.1% for the embryo test, 7.50*×*10^*−*5^*U. nuda/H. vulgare* DNA copies for the qPCR method, and 35 infected ears per 100 *m*^2^ for the field observation.

We constructed confusion matrices to compare the actual values from the binary classifications of the field results to the predicted values from the binary classifications of each laboratory detection method (Supplementary Information, Table 7). Samples for which the laboratory detection method correctly classified as above the threshold (based on the reference field results), were assigned as true positives (TP), while samples that both methods classified as below the threshold were considered true negatives (TN). False negatives (FN) were assigned to cases with infections above the threshold in the field but incorrectly measured as below the threshold according to the respective laboratory detection method. Conversely, samples that were below the tolerance threshold in the field but were classified as above the threshold by the respective laboratory detection method were considered false positives (FP).

Each laboratory detection method’s confusion matrix was used to calculate performance metrics —sensitivity, specificity, precision, the false discovery rate, and accuracy —to directly compare the qPCR method and the embryo test. Sensitivity and specificity measure the ability of the laboratory detection methods to correctly classify the samples as above or below the tolerance thresholds, respectively, while precision and false discovery rate evaluate the reliability of the laboratory detection methods’ predicted value. Accuracy, on the other hand, represents the overall proportion of the correctly classified samples by the laboratory detection methods. We calculated these performance metrics with the following formulas:

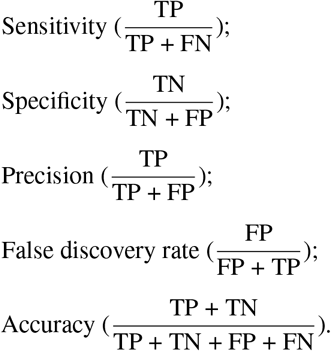

## Data Availability

All data collected from the experiments and used in the analyses are available in the Supplementary Information or the provided data tables.

## Acknowledgements

We thank Nicole Bischofberger and Damian Amrein for assistance in acquiring seed material, and Francesco Bassi and Magnus Wagner for help with sample processing. We are grateful to Prof. Gunther Doehlemann (University of Cologne) for *Ustilago hordei* and *Ustilago maydis* strains, and Marco Wüthrich for providing cultures of *Tilletia caries, Tilletia controversa, Ustilago* spp., *Tilletiopsis* spp., *Pseudozyma* spp., *Entyloma* spp., and *Holtermaniella* spp. We acknowledge Matteo Selmi for providing uninfected barley seeds from the “Mixture Ceccarelli” unknown cultivar, grown in Loritto, Italy. We thank Daniel Fuchs and his team for assistance with field trials and German Bonilla-Rosso for statistical discussions. We thank Joëlle Schläpfer for her comments on the manuscript, and we appreciate the early discussions with Annette Büttner-Mainik.

## Funding

This research was supported by Fondation Sur-la-Croix, IP Suisse, the Swiss Association of Cereal Producers, Swiss granum, and swisssem.

## Author contributions statement

Conceptualization: C.P. and K.E.S; Formal analysis: C.P. and K.E.S; Funding acquisition: K.E.S and S.V.; Investigation: C.P., E.J., I.B., P.B., A.K., and K.E.S.; Methodology: C.P., E.J., and K.E.S; Project administration K.E.S.; Resources: S.V., T.H., and F.W.; Supervision: K.E.S. and D.C.; Visualization: C.P. and K.E.S.; Writing-original draft preparation: C.P. with support from K.E.S; Writing-review & editing: C.P., K.E.S., D.C., S.V., T.H., F.W., P.B., E.J., I.B., and A.K.

## Additional information

### Competing interests

The authors declare no competing financial and/or non-financial interests in relation to the work described.

## Notes

### Competing Interest Statement

The authors have declared no competing interest.

