## Supplemental Information for "A new molecular seed assay to predict *Ustilago nuda* field infection levels"

### Supplementary Figures

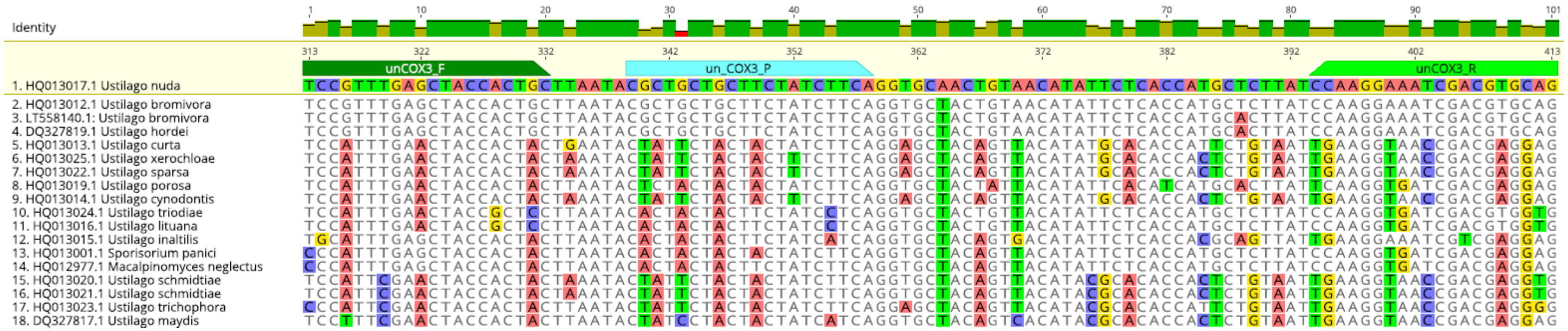


**Supplementary Figure 1. Alignment of COX3 target gene sequences**. The GenBank accession numbers for sequences included in the alignments are shown in sequence names, including the *Ustilago nuda* reference sequence, HQ013017.1. We included closely related *Ustilago* species and additional fungi species that may be amplified based on the Primer BLAST program. Primers (unCOX3_F and unCOX3_R) are highlighted in two green colors, while the probe is highlighted in light blue (unCOX3_P).

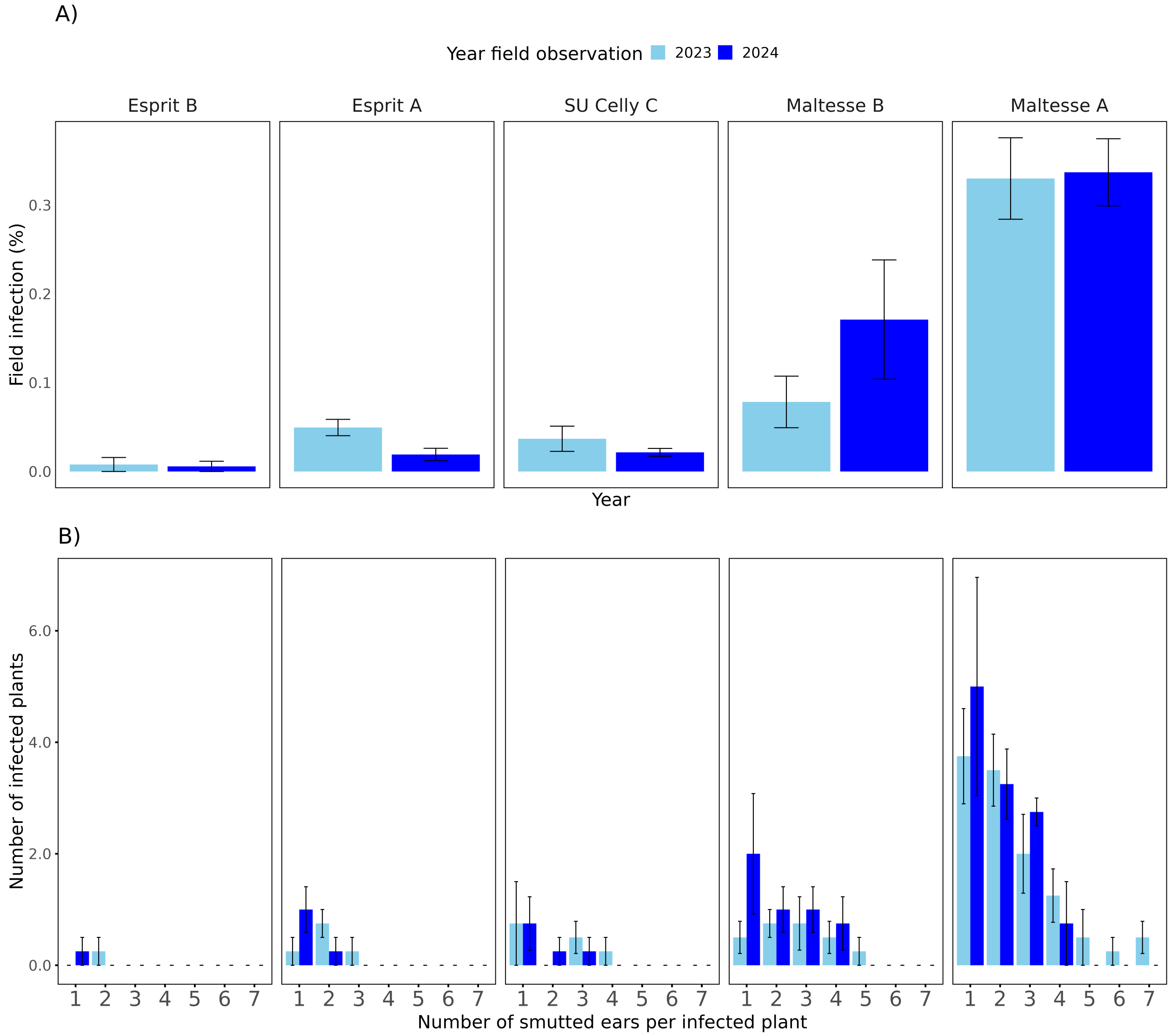


**Supplementary Figure 2.** ***Ustilago nuda* field infection levels and number of smutted ears per infected plant.** Five of the six commercially available seed lots showed smutted ears. The number of smutted ears per plant was evaluated across two years of field observations: 2023 (light blue) and 2024 (dark blue). (A) Field infection levels from each seed lot, and (B) the distribution of smutted ears per infected plant. In both panels, error bars represent the standard error. The sixth commercially available seed lot, KWS Orbit, had no observed field infections.

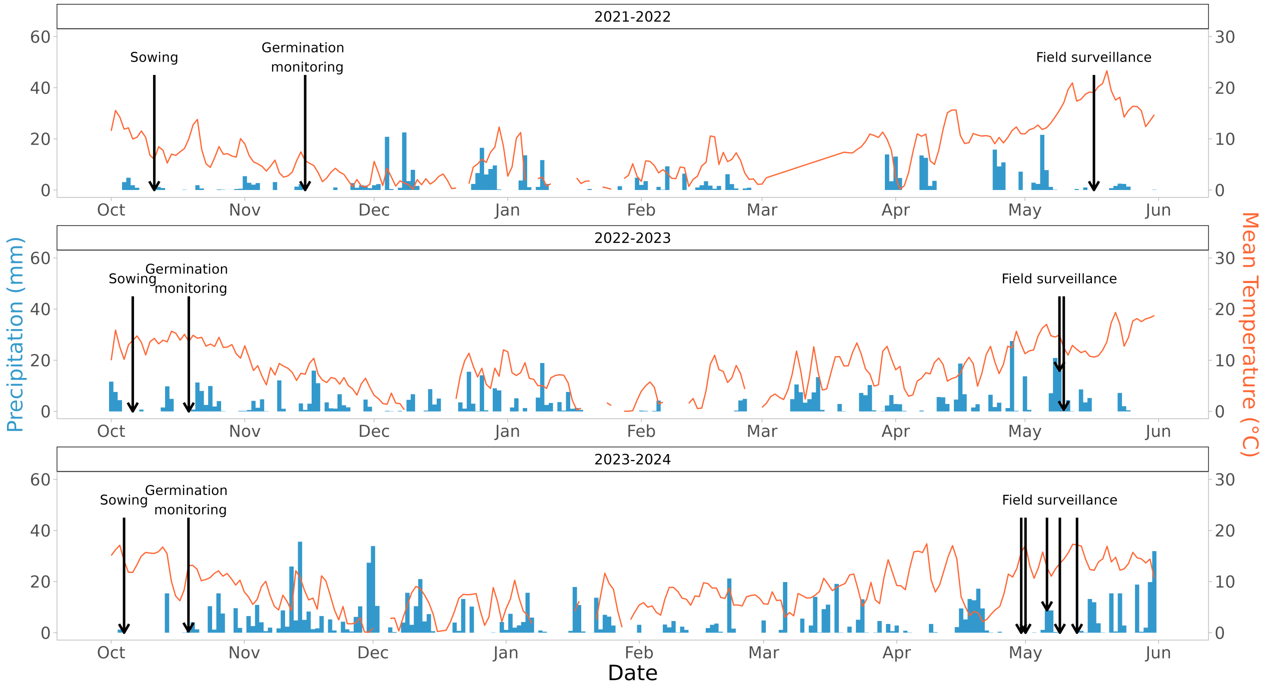


**Supplementary Figure 3. Weather data collected during field trials.** The weather measurements were taken at the MeteoSwiss station Zurich/Affoltern, which was closest to the fields. Daily mean temperature (°C) is shown as an orange line and total precipitation (mm) per day as blue bars. Arrows indicate the dates of seed sowing, germination monitoring, and field surveillance.

### Supplementary Tables

| Source | Infection status | Concentration (ng/μL)[a] | Newly developed COX3 TaqMan protocol | | | Published ITS SYBR green protocol [b] | | |
|---|---|---|---|---|---|---|---|---|
| | | | Cq Mean[c] | Cq Std. Dev [d] | CV (%) [e] | Cq Mean | Cq Std. Dev | CV (%) |
| ***Hordeum vulgare* seedling** | *Ustilago nuda* infected | 5.00 | 25.43 | 0.11 | 0.44 | 27.92 | 1.13 | 4.05 |
| | Uninfected | | - | - | - | - | - | - |
| ***Hordeum vulgare* seed** | *Ustilago nuda* infected | 5.00 | 29.74 | 0.09 | 0.29 | 27.97 | 0.74 | 2.66 |
| | Uninfected | | - | - | - | - | - | - |

**Supplementary Table 1. Comparison of relative variance in *Ustilago nuda* amplification with our newly designed COX3 primers in a Taqman protocol and the published ITS primers in a SYBR green protocol.** Our COX3 primers were included in a multiplex reaction, and the published ITS primers were run in a singleplex reaction to detect *U. nuda*. A difference between the coefficients of variation (CV) were found between the tested protocols. A sample each of infected and uninfected *Hordeum vulgare* seeds and seedlings was used. Seed DNA was extracted from a 0.02 g flour subsample of 2000 milled seeds. The seeds were of the cultivar KWS Orbit; the seed lot was not otherwise included in this study. Seedlings were grown from a SU Celly seed lot used in this study, which had been treated with warm water to obtain an uninfected sample. The tested seedling DNA samples were extracted from a pool sample of 20 seedling segments (0.5 cm each) cut from the base of seedlings grown in soil for 5 days (uninfected seedlings) and 14 days (infected seedlings). Each sample was analyzed with triplicate reactions using the same template DNA concentration for both protocols. All uninfected samples that showed no amplification and are indicated by the symbol '-'.

[a]Final DNA concentration in each qPCR reaction
[b]SYBR Green Protocol with ITS primers published in a previous study[36]
[c]Average cycle of quantification across replicates
[d]Standard deviation of the cycle of quantification among replicates
[e]Relative variance or coefficient of variation ($^{\text{Cq Std. Dev.}}/_{\text{Cq Mean}}$)*100

| Low-infected seed lot (%) | High-infected seed lot (%) | Cultivars | | |
|---|---|---|---|---|
| | | Azrah | SU Celly | Semper |
| | | ($\overline{Cq}\ \overline{Mean}$ ± Std. Dev.[a]) | | |
| 100 | 0 | 24.26 ± 0.52 | 25.48 ± 0.16 | 24.42 ± 0.42 |
| 99 | 1 | 24.36 ± 0.66 | 25.63 ± 0.24 | 24.53 ± 0.45 |
| 90 | 10 | 24.62 ± 0.17 | 25.53 ± 0.25 | 24.38 ± 0.43 |
| 0 | 100 | 24.39 ± 0.46 | 25.61 ± 0.28 | 24.50 ± 0.36 |

**Supplementary Table 2. Evaluation of *Hordeum vulgare* GAPDH target stability in three cultivars with different *Ustilago nuda* infection levels.** This evaluation is based on HEX fluorescence measurements targeting *H. vulgare* GAPDH and are reported as the average of the mean quantification cycle (Cq Mean) ± standard deviation. Average Cq Mean values were calculated for high- and low-infected seed lots from three cultivars and their mixtures. These seed lots and mixtures are part of an experiment designed to assess the relationship between *U. nuda* infection levels observed in the field and those detected using laboratory methods (Figure 2).

[a]Average of the mean quantification cycle (Cq) values across tested seed lots ± standard deviation among the seed lot subsample Cq Mean

| Seed number tested | Flour amount for DNA extraction (g) | Number of flour batches tested | Cultivars | | | | | |
|---|---|---|---|---|---|---|---|---|
| | | | Esprit A | KWS Orbit | Maltesse A | Esprit B | SU Celly C | Maltesse B |
| | | | ($\overline{Cq}\ \overline{Mean}$ ± Std. Dev.[a]) | | | | | |
| 2000 | 0.02 | 1 | 24.70 ± 0.17 | 24.66 ± 0.52 | 25.74 ± 0.52 | 24.68 ± 0.38 | 25.49 ± 0.21 | 25.75 ± 0.38 |
| | | 10 | 24.50 ± 0.18 | 24.64 ± 0.32 | 25.42 ± 0.21 | | | |
| | 0.2 | | 24.70 ± 0.31 | 24.60 ± 0.34 | 25.63 ± 0.45 | | | |
| 7000 | 0.02 | | 24.58 ± 0.30 | 24.63 ± 0.33 | 25.65 ± 0.39 | | | |

**Supplementary Table 3. Evaluation of *Hordeum vulgare* GAPDH target stability in six commercial seed lots.** This evaluation is based on HEX fluorescence measurements targeting *H. vulgare* GAPDH and the results are reported as the average of the mean quantification cycle (Cq Mean) ± standard deviation. Average Cq Mean values were calculated for six commercial *H. vulgare* seed lots. These seed lots were used for the comparison of *U. nuda* infection obeserved in the field to those measured using laboratory detection method 3. In this comparison, each seed lot was represented by 10 technical replicates from 1 milled flour batch of 2000 seeds, and 0.02 g of flour used for each DNA extraction. Three of the six seed lots (Espirit A, KWS Orbit, and Maltesse A) were additionally used in an experiment to test key parameters for establishing a representative *H. vulgare* seed lot samples for *Ustilago nuda* qPCR detection (Figure 1). For these three seed lots, the average Cq Mean values are reported for each tested parameter level, which included different seed number (2000 and 7000 seeds), flour amount for DNA extraction (0.02 and 0.2 g),the number of flour batches used for ten DNA sub-sample extraction (one batch extracted ten times or ten batches extracted one time each).

[a]Average of the mean quantification cycle (Cq) values across tested seed lots ± standard deviation among the seed lot subsample Cq Mean

| Seed lot | Key parameter | Parameter level | Mean measured infection ($U. nuda$ / $H. vulgare$ DNA copies) | Min measured infection ($U. nuda$ / $H. vulgare$ DNA copies) | Max measured infection ($U. nuda$ / $H. vulgare$ DNA copies) | Median measured infection ($U. nuda$ / $H. vulgare$ DNA copies) | Std. Error | Std. Deviation | CV (%)[1] | Kruskal test $\chi^2$ | Kruskal test p-value |
|---|---|---|---|---|---|---|---|---|---|---|---|
| Esprit A | Seed number tested[2] | 2000 seed | $4.8 \times 10^{-4}$ | 0.0 | $3.1 \times 10^{-3}$ | 0.0 | $3.4 \times 10^{-4}$ | $1.1 \times 10^{-3}$ | 221.3 | 0.53 | 0.47 |
| | | 7000 seed | $8.2 \times 10^{-5}$ | 0.0 | $8.2 \times 10^{-4}$ | 0.0 | $8.2 \times 10^{-5}$ | $2.6 \times 10^{-4}$ | 221.3 | | |
| | Flour amount for DNA extraction[3] | 0.02 g flour | $4.8 \times 10^{-4}$ | 0.0 | $3.1 \times 10^{-3}$ | 0.0 | $3.4 \times 10^{-4}$ | $1.1 \times 10^{-3}$ | 221.3 | 0.31 | 0.58 |
| | | 0.2 g flour | $3.5 \times 10^{-4}$ | 0.0 | $1.7 \times 10^{-3}$ | 0.0 | $1.8 \times 10^{-4}$ | $5.6 \times 10^{-4}$ | 159.2 | | |
| | Flour batch number for DNA extraction[4] | 1 batch | 0.0 | 0.0 | 0.0 | 0.0 | 0.0 | 0.0 | - | 2.11 | 0.15 |
| | | 10 batches | $4.8 \times 10^{-4}$ | 0.0 | $3.1 \times 10^{-3}$ | 0.0 | $3.4 \times 10^{-4}$ | $1.1 \times 10^{-3}$ | 221.3 | | |
| KWS Orbit | Seed number tested | 2000 seed | 0.0 | 0.0 | 0.0 | 0.0 | 0.0 | 0.0 | - | - | - |
| | | 7000 seed | 0.0 | 0.0 | 0.0 | 0.0 | 0.0 | 0.0 | - | | |
| | Flour amount for DNA extraction | 0.02 g flour | 0.0 | 0.0 | 0.0 | 0.0 | 0.0 | 0.0 | - | 1.00 | 0.32 |
| | | 0.2 g flour | $8.7 \times 10^{-5}$ | 0.0 | $7.1 \times 10^{-4}$ | 0.0 | $7.1 \times 10^{-5}$ | $2.2 \times 10^{-4}$ | 256.4 | | |
| | Flour batch number for DNA extraction | 1 batch | 0.0 | 0.0 | 0.0 | 0.0 | 0.0 | 0.0 | - | - | - |
| | | 10 batches | 0.0 | 0.0 | 0.0 | 0.0 | 0.0 | 0.0 | - | | |
| Maltesse A | Seed number tested | 2000 seed | $5.8 \times 10^{-3}$ | $9.3 \times 10^{-4}$ | $1.3 \times 10^{-2}$ | $5.2 \times 10^{-3}$ | $1.4 \times 10^{-3}$ | $4.5 \times 10^{-3}$ | 76.5 | 0.21 | 0.65 |
| | | 7000 seed | $6.5 \times 10^{-3}$ | $1.3 \times 10^{-3}$ | $1.3 \times 10^{-2}$ | $6.7 \times 10^{-3}$ | $1.4 \times 10^{-3}$ | $4.4 \times 10^{-3}$ | 68.5 | | |
| | Flour amount for DNA extraction | 0.02 g flour | $5.8 \times 10^{-3}$ | $9.3 \times 10^{-4}$ | $1.3 \times 10^{-2}$ | $5.2 \times 10^{-3}$ | $1.4 \times 10^{-3}$ | $4.5 \times 10^{-3}$ | 76.5 | 1.120 | 0.290 |
| | | 0.2 g flour | $8.0 \times 10^{-3}$ | $2.7 \times 10^{-3}$ | $1.9 \times 10^{-2}$ | $6.0 \times 10^{-3}$ | $1.6 \times 10^{-3}$ | $5.1 \times 10^{-3}$ | 63.8 | | |
| | Flour batch number for DNA extraction | 1 batch | $1.2 \times 10^{-2}$ | $4.3 \times 10^{-3}$ | $4.3 \times 10^{-2}$ | $7.9 \times 10^{-3}$ | $3.7 \times 10^{-3}$ | $1.2 \times 10^{-2}$ | 95.1 | 2.64 | 0.10 |
| | | 10 batches | $5.8 \times 10^{-3}$ | $9.3 \times 10^{-4}$ | $1.3 \times 10^{-2}$ | $5.2 \times 10^{-3}$ | $1.4 \times 10^{-3}$ | $4.5 \times 10^{-3}$ | 76.5 | | |

**Supplementary Table 4. Evaluation of key parameters to establish a representative *Hordeum vulgare* seed lot sample for *Ustilago nuda* qPCR detection.** The evaluated key parameters included: the number of seeds tested (2000 versus 7000), the amount of flour used for the DNA extraction (0.02 g versus 0.2 g), and the number of flour batches used for the ten DNA sub-sample extractions (one batch extracted ten times versus ten batches extracted one time each). The later analysis was used to assess inter- and intra-flour batch *U. nuda* DNA variation. The key parameters were tested with three different seed lots, which each consist in a different cultivar. The mean, minimum, maximum, and median measured infections detected in all parameter level are reported as the ratio $U. nuda / H. vulgare$ DNA copies. For each parameter level, the standard errors, standard deviation, coefficient of variation, and the Kruskal-Wallis test statistics ($\chi^2$ and p-values) are reported. The Kruskal test p-values were used to evaluate whether differences between parameter levels were significantly different within cultivars. Coefficients of variation and Kruskal test results that were not possible to calculate due to zero values are indicated using the symbol "-".

[1] Coefficient of variation ($\text{Std. Dev. infected embryo observed} / \text{Infected embryo observed Mean}$)*100

[2] For the key parameter of seed number tested, 10 separate flour batches were used for the DNA extractions of 0.02 g of flour.

[3] For the key parameter of flour amount, 10 separate flour batches were used for the DNA extractions of 2000 seeds.

[4] For the key parameter of flour batch number for DNA extraction, 0.02 g of flour milled from 2000 seeds were used.

| Tested model | Lab method | $R^2$ | $R^2$ adj | MSE[a] | AIC[b] | BIC[c] |
|---|---|---|---|---|---|---|
| Linear | qPCR | 0.974 | 0.972 | $2.69 \times 10^{-6}$ | -113.860 | -112.405 |
| | Embryo | 0.756 | 0.732 | 0.420 | 29.645 | 31.099 |
| Polynomial | qPCR | 0.974 | 0.969 | $2.69 \times 10^{-6}$ | -111.867 | -109.927 |
| | Embryo | 0.844 | 0.810 | 0.268 | 26.253 | 28.193 |
| Exponential | qPCR | -1.677 | - | $2.80 \times 10^{-4}$ | 40.129 | 41.584 |
| | Embryo | 0.791 | - | 0.361 | 31.767 | 32.675 |

**Supplementary Table 5. Model evaluations used to derive the tolerance threshold for the qPCR method.** The models are based on the correlation between *Ustilago nuda* infection detected with the qPCR method and field observations. The linear model was chosen for its predictive performance (highest adjusted $R^2$) and lower complexity (lowest AIC and BIC) compared to the polynomial and exponential models.

[a] Mean squared error
[b] Akaike information criterion
[c] Bayesian information criterion

| Seed lot | Plot repetition | Year collecting data | Infection observed (infected ears per 100 $m^2$) | Tolerance threshold (infected ears per 100 $m^2$) | Plot infection compared to the tolerance threshold | Average infection (infected ears per 100 $m^2$) | Average infection compared to the tolerance threshold |
|---|---|---|---|---|---|---|---|
| KWS Orbit | 1 | 2023 | 0 | 35 | Below | 0 | Below |
| | 2 | | 0 | | Below | | |
| | 3 | | 0 | | Below | | |
| | 4 | | 0 | | Below | | |
| | 5 | 2024 | 0 | | Below | | |
| | 6 | | 0 | | Below | | |
| | 7 | | 0 | | Below | | |
| | 8 | | 0 | | Below | | |
| Esprit B | 1 | 2023 | 0 | 35 | Below | 2 | Below |
| | 2 | | 0 | | Below | | |
| | 3 | | 0 | | Below | | |
| | 4 | | 10 | | Below | | |
| | 5 | 2024 | 5 | | Below | | |
| | 6 | | 0 | | Below | | |
| | 7 | | 0 | | Below | | |
| | 8 | | 0 | | Below | | |
| Esprit A | 1 | 2023 | 16 | 35 | Below | 10 | Below |
| | 2 | | 10 | | Below | | |
| | 3 | | 10 | | Below | | |
| | 4 | | 15 | | Below | | |
| | 5 | 2024 | 16 | | Below | | |
| | 6 | | 5 | | Below | | |
| | 7 | | 0 | | Below | | |
| | 8 | | 10 | | Below | | |
| SU Celly C | 1 | 2023 | 0 | 35 | Below | 11 | Below |
| | 2 | | 16 | | Below | | |
| | 3 | | 15 | | Below | | |
| | 4 | | 21 | | Below | | |
| | 5 | 2024 | 10 | | Below | | |
| | 6 | | 10 | | Below | | |
| | 7 | | 0 | | Below | | |
| | 8 | | 15 | | Below | | |
| Maltesse B | 1 | 2023 | 36 | 35 | Above | 43 | Above |
| | 2 | | 10 | | Below | | |
| | 3 | | 77 | | Above | | |
| | 4 | | 15 | | Below | | |
| | 5 | 2024 | 92 | | Above | | |
| | 6 | | 51 | | Above | | |
| | 7 | | 56 | | Above | | |
| | 8 | | 5 | | Below | | |
| Maltesse A | 1 | 2023 | 83 | 35 | Above | 134 | Above |
| | 2 | | 203 | | Above | | |
| | 3 | | 143 | | Above | | |
| | 4 | | 164 | | Above | | |
| | 5 | 2024 | 124 | | Above | | |
| | 6 | | 104 | | Above | | |
| | 7 | | 158 | | Above | | |
| | 8 | | 98 | | Above | | |

**Supplementary Table 6. Consistency of *Ustilago nuda* observed infections in individual field plots compared to the field tolerance threshold.** For six commercially available seed lots, the number of infected ears per 100 $m^2$ were assessed over two years in four field plot each year. The Infection level observed in each plot and their averages were classified as either above or below the field tolerance threshold.

A)

| Predicted values (qPCR method) | | Actual values (Field reference): Above tolerance threshold | Actual values (Field reference): Below tolerance threshold |
|---|---|---|---|
| | Above tolerance threshold | True Positive (TP):<br>- Maltesse A<br>- Maltesse B | False Positive (FP):<br>— |
| | Below tolerance threshold | False Negative (FN):<br>— | True negative (TN):<br>- Esprit A<br>- Esprit B<br>- SU Celly C<br>- KWS Orbit |

B)

| Actual values (Field reference): Above tolerance threshold | Actual values (Field reference): Below tolerance threshold | | Predicted values (Embryo Test) |
|---|---|---|---|
| True Positive (TP):<br>— | False Positive (FP):<br>— | Above tolerance threshold | |
| False Negative (FN):<br>- Maltesse A<br>- Maltesse B | True negative (TN):<br>- Esprit A<br>- Esprit B<br>- SU Celly C<br>- KWS Orbit | Below tolerance threshold | |

**Supplementary Table 7. Confusion matrices comparing the predicted values of the qPCR method and the embryo test with the actual values of the field observations.** For the confusion matrix based on the (A) qPCR method, the predicted values of the *Ustilago nuda* infection levels were classified as either above or below the qPCR tolerance threshold: $7.50 \times 10^{-5}$ *U. nuda*/*H. vulgare* DNA copies. For the confusion matrix based on the (B) embryo test, predicted values were classified as either above or below the tolerance threshold of 0.1% infected embryos. Seed lots were classified as either above or below the field tolerance threshold of 35 infected ears per 100 $m^2$. based on the number of observed ear infections in the field. These classifications served as actual values in the confusion matrices. The predicted values that matched the actual values were either true positive or true negative, while the predicted values that did not match the actual values were either false positive or false negative.

| Cultivar | Low-infected seed lot (%) | High-infected seed lot (%) | Evaluator | Infected embryos observed | CV(%)[a] |
|---|---|---|---|---|---|
| Azrah | 100 | 0 | 1 | 20 | 29.4 |
| | | | 2 | 22 | |
| | | | 3 | 12 | |
| | 99 | 1 | 1 | 28 | 21.6 |
| | | | 2 | 24 | |
| | | | 3 | 18 | |
| | 90 | 10 | 1 | 6 | 50.0 |
| | | | 2 | 6 | |
| | | | 3 | 2 | |
| | 0 | 100 | 1 | 22 | 20.4 |
| | | | 2 | 32 | |
| | | | 3 | 24 | |
| Semper | 100 | 0 | 1 | 10 | 63.0 |
| | | | 2 | 10 | |
| | | | 3 | 2 | |
| | 99 | 1 | 1 | 14 | 70.5 |
| | | | 2 | 10 | |
| | | | 3 | 2 | |
| | 90 | 10 | 1 | 40 | 28.8 |
| | | | 2 | 32 | |
| | | | 3 | 22 | |
| | 0 | 100 | 1 | 120 | 20.4 |
| | | | 2 | 96 | |
| | | | 3 | 80 | |

**Supplementary Table 8. Inter-evaluator result differences in visual embryo test.** Discrepancy in the number of *Ustilago nuda* infected embryos detected in eight seed samples by three independent evaluators with the validated International Seed Test Association test[24]. The seed samples were composed of 1000 embryos extracted from two different cultivars (Azrah and Semper), including low- and high-infected seed lots and their mixtures. The variability in the number of infected embryos among evaluators and seed samples was assessed using the coefficient of variation.

[a]Coefficient of variation $(\text{Std. Dev. infected embryo observed}/\text{Infected embryo observed Mean}) * 100$

| Seed lot | Sub-sample repetition | Measured infection (*U. nuda*/*H. vulgare* DNA copies) | Tolerance threshold (*U. nuda*/*H. vulgare* DNA copies) | Sub-sample infection compared to the tolerance threshold | Average infection (*U. nuda*/*H. vulgare* DNA copies) | Average infection compared to the tolerance threshold |
|---|---|---|---|---|---|---|
| KWS Orbit | 1 | 0.0 | $7.5 \times 10^{-5}$ | Below | 0.0 | Below |
| | 2 | 0.0 | | Below | | |
| | 3 | 0.0 | | Below | | |
| | 4 | 0.0 | | Below | | |
| | 5 | 0.0 | | Below | | |
| | 6 | 0.0 | | Below | | |
| | 7 | 0.0 | | Below | | |
| | 8 | 0.0 | | Below | | |
| | 9 | 0.0 | | Below | | |
| | 10 | 0.0 | | Below | | |
| Esprit B | 1 | 0.0 | $7.5 \times 10^{-5}$ | Below | 0.0 | Below |
| | 2 | 0.0 | | Below | | |
| | 3 | 0.0 | | Below | | |
| | 4 | 0.0 | | Below | | |
| | 5 | 0.0 | | Below | | |
| | 6 | 0.0 | | Below | | |
| | 7 | 0.0 | | Below | | |
| | 8 | 0.0 | | Below | | |
| | 9 | 0.0 | | Below | | |
| | 10 | 0.0 | | Below | | |
| Esprit A | 1 | 0.0 | $7.5 \times 10^{-5}$ | Below | 0.0 | Below |
| | 2 | 0.0 | | Below | | |
| | 3 | 0.0 | | Below | | |
| | 4 | 0.0 | | Below | | |
| | 5 | 0.0 | | Below | | |
| | 6 | 0.0 | | Below | | |
| | 7 | 0.0 | | Below | | |
| | 8 | 0.0 | | Below | | |
| | 9 | 0.0 | | Below | | |
| | 10 | 0.0 | | Below | | |
| SU Celly C | 1 | 0.0 | $7.5 \times 10^{-5}$ | Below | 0.0 | Below |
| | 2 | 0.0 | | Below | | |
| | 3 | 0.0 | | Below | | |
| | 4 | 0.0 | | Below | | |
| | 5 | 0.0 | | Below | | |
| | 6 | 0.0 | | Below | | |
| | 7 | 0.0 | | Below | | |
| | 8 | 0.0 | | Below | | |
| | 9 | 0.0 | | Below | | |
| | 10 | 0.0 | | Below | | |
| Maltesse B | 1 | 0.0 | $7.5 \times 10^{-5}$ | Below | $9.1 \times 10^{-4}$ | Above |
| | 2 | $1.3 \times 10^{-3}$ | | Above | | |
| | 3 | $6.7 \times 10^{-4}$ | | Above | | |
| | 4 | $1.1 \times 10^{-3}$ | | Above | | |
| | 5 | $1.2 \times 10^{-3}$ | | Above | | |
| | 6 | $9.1 \times 10^{-4}$ | | Above | | |
| | 7 | $1.6 \times 10^{-3}$ | | Above | | |
| | 8 | 0.0 | | Below | | |
| | 9 | $8.1 \times 10^{-4}$ | | Above | | |
| | 10 | $1.5 \times 10^{-3}$ | | Above | | |
| Maltesse A | 1 | $1.5 \times 10^{-2}$ | $7.5 \times 10^{-5}$ | Above | $1.2 \times 10^{-2}$ | Above |
| | 2 | $5.4 \times 10^{-3}$ | | Above | | |
| | 3 | $9.5 \times 10^{-3}$ | | Above | | |
| | 4 | $8.2 \times 10^{-3}$ | | Above | | |
| | 5 | $4.3 \times 10^{-2}$ | | Above | | |
| | 6 | $4.3 \times 10^{-3}$ | | Above | | |
| | 7 | $1.7 \times 10^{-2}$ | | Above | | |
| | 8 | $6.9 \times 10^{-3}$ | | Above | | |
| | 9 | $7.6 \times 10^{-3}$ | | Above | | |
| | 10 | $7.8 \times 10^{-3}$ | | Above | | |

**Supplementary Table 9. Consistency of *Ustilago nuda* infections detected with qPCR in individual DNA extractions compared to the qPCR tolerance threshold.** For six commercially available seed lots, the measured infections were assessed with the ratio *U. nuda*/*H. vulgare* DNA copies in ten DNA extractions. The Infection level measured in each DNA extraction and their averages were classified as either above or below the qPCR tolerance threshold.

| Template DNA | Template DNA concentration (ng/µL)[5] | Test | Spiked gBlock gene fragments | Spiked gBlock gene copy numbers | *Ustilago nuda* COX3 | | *Hordeum vulgare* GADPH | |
|---|---|---|---|---|---|---|---|---|
| | | | | | Cq Mean | Cq Std. Dev. | Cq Mean | Cq Std. Dev. |
| *H. vulgare* seed | 5.0 | qPCR inhibition | UnCOX3_stdcurve | 62500 | 21.15 | 0.06 | 24.79 | 0.46 |
| | 2.5 | | | | 21.23 | 0.18 | 26.04 | 0.63 |
| | 0.8 | | | | 21.43 | 0.32 | 28.1 | 0.48 |
| | 5.0 | Control | - | - | 0.00 | 0.00 | 25.11 | 0.10 |
| *H. vulgare* seedling | 5.0 | qPCR inhibition | UnCOX3_stdcurve | 62500 | 21.16 | 0.13 | 24.20 | 0.28 |
| | 2.5 | | | | 21.33 | 0.04 | 25.42 | 0.36 |
| | 0.8 | | | | 21.29 | 0.10 | 28.94 | 0.27 |
| | 2.5 | Control | - | - | 0.00 | 0.00 | 25.78 | 0.15 |
| *Lens culinaris* seed | 5.0 | qPCR inhibition | UnHvGADPH_stdcurve | 1563000 | 0.00 | 0.00 | 17.51 | 0.12 |
| | 2.5 | | | | 0.00 | 0.00 | 17.58 | 0.21 |
| | 0.8 | | | | 0.00 | 0.00 | 17.45 | 0.18 |
| | 5.0 | Control | - | - | 0.00 | 0.00 | 0.00 | 0.00 |

**Supplementary Table 10. Evaluation of potential qPCR inhibition.** DNA from *Hordeum vulgare* seeds and seedlings, and *Lens culinaris* seeds were spiked with gBlock gene fragments. Uninfected *H. vulgare* seeds were obtained from an undefined cultivar "mixture Ceccarelli" grown in Loritto, Italy. DNA was extracted from a 0.02 g flour subsample of 2000 milled seeds. *H. vulgare* seedlings were grown in soil for 5 days from a seed lot of cultivar SU Celly that had been treated with warm water. DNA was extracted from a pooled subsample of 20 seedling segments (0.5 cm each). *L. culinaris* DNA was extracted from seeds intended for consumption.

| Plant | Isolation sources | Closest accession and region | Database | Percent similarity (%) | Query cover (%) | Classification of closest BLAST result |
|---|---|---|---|---|---|---|
| *Triticum aestivum* | Seed | MN901739: 352-1175 | Core nucleotide database (core_nt) | 95.3 | 100 | *Entyloma* spp. |
| *Hordeum vulgare* | Smutted head | JALCCE020000007:289349-290325 | Whole genome shotgut contigs (Basidiomycetes) | 100 | 100 | *Pseudozyma flocculosa* |
| *Triticum aestivum* | seed | MW248454: 151-1076 | Core nucleotide database (core_nt) | 100 | 100 | *Holtermanniella festucosa* |
| *Hordeum vulgare* | Smutted ear | DQ835992: 75-563 | Core nucleotide database (core_nt) | 96.33 | 100 | *Tilletiopsis minor* |
| *Glycine max* | seed | LVYE01000169:3537-4519 | Whole genome shotgut contigs (Basidiomycetes) | 100 | 100 | *Ustilago trichophora* |

**Supplementary Table 11. Information about fungal isolates from Swiss cereal samples.** The fungi were isolated from either seeds or smutted ears and cultured on PDA plates to obtain mycelia. DNA was extracted from each isolate's mycelia and used to test the *Ustilago nuda* primer specificity (Table 1).

| Gene target | Target organism | Oligo name | Sequence (5' - 3') | Length (bp) | Genbank accession number |
|---|---|---|---|---|---|
| COX3 | *Ustilago nuda* | UnCOX3_stdcurve | TCC GTT TGA GCT ACC ACT GCT<br>TAA TAC GCT GCT GCT TCT ATC<br>TTC AGG TGC AAC TGT AAC ATA<br>TTC TCA CCA TGC TCT TAT CCA AGG<br>AAA TCG ACG TGC AG | 125 | HQ013017 |
| GADPH | *Hordeum vulgare* | HvGADPH_stdcurve | CAG TTC ACG GCC ATT GGA AGC<br>ACA GTG ACA TCA AGC TCA AAG<br>ACG ACA AGA CGC TGC TCT TCG<br>GCG AGA A G CCA GTT ACT GTC<br>TTT GGC GTC AGG TAG TAT TGA T | 125 | MT933276 |

**Supplementary Table 12. Sequences of gBlock gene fragments used as the standard curve.** The standard curves were included in all qPCR plates for the absolute quantification of *Ustilago nuda* and *Hordeum vulgare*. The copy numbers of each target were calculated from the absolute quantification and used to normalize the pathogen DNA to its host's DNA. The GenBank accession numbers identify the sequences used as the basis of the gBlock gene fragments.

### Supplementary Methods

The fungal isolates provided by Marco Wüthrich were isolated from Swiss cereal, either from seeds or smutted ears. These fungal isolates were grown on PDA plates to obtain mycelia from which DNA was extracted with the NucleoSpin Plant II Kit (MACHEREY-NAGEL, Germany). The extracted DNA was amplified using the primers ITS1 (5'-TCC GTA GGT GAA CCT GCG G-3') and LR5 (5'-TCCTGAGGGAAACTTCG-3'). Each PCR reaction had a total volume of 20 $\mu$l, containing 1× GoTaq Flexi Colorless Buffer, 0.4 $\mu$M of each primer, 2.5 $\mu$M $MgCl_2$, 0.2 mM dNTPs, and 0.05 U/$\mu$l GoTaq G2 Hot Start Polymerase (Promega, USA). A total of 15 ng of DNA template was used per reaction. Sanger sequences was performed by Microsynth (Balgach, Switzerland) using the ITS1 primer. Low quality regions at the sequence end were trimmed until the base pair quality scores were greater than 40. Sequences were then searched using BLAST against both the core nucleotide database (core_nt) and the whole-genome shotgun contigs (wgs) databases restricted to Basidiomycetes. The closest matches are reported and were used to deduce the isolate's identity that were used for primer specificity testing. Sequences will be deposited in NCBI (accession numbers of sequences to be provided prior to acceptance).
